# Enhanced cell-cell contact stability and decreased N-cadherin-mediated migration upon Fibroblast Growth Factor Receptor-N-cadherin cross-talk

**DOI:** 10.1101/465930

**Authors:** Thao Nguyen, Laurence Duchesne, Gautham Hari Narayana Sankara Narayana, Nicole Boggetto, David D. Fernig, Chandrashekhar Uttamrao Murade, Benoit Ladoux, René-Marc Mège

## Abstract

N-cadherin adhesion has been reported to enhance cancer and neuronal cell migration either by mediating actomyosin-based force transduction or initiating Fibroblast Growth Factor Receptor (FGFR)-dependent biochemical signalling. Here we show that FGFR1 reduces N-cadherin-mediated cell migration. Both proteins are co-stabilised at cell-cell contacts through direct interaction. As a consequence, cell adhesion is strengthened, limiting the migration of cells on N-cadherin. Both the inhibition of migration and the stabilisation of cell adhesions require the FGFR activity stimulated by N-cadherin engagement. FGFR1 stabilises N-cadherin at the cell membrane through a pathway involving Src and p120. Moreover, FGFR1 stimulates the anchoring of N-cadherin to actin. We found that the migratory behaviour of cells depends on an optimum balance between FGFR-regulated N-cadherin adhesion and actin dynamics. Based on these findings we propose a positive feedback loop between N-cadherin and FGFR at adhesion sites limiting N-cadherin-based single cell migration.

## Introduction

Cell adhesion and migration are central processes in morphogenesis, wound healing and cancerogenesis. Cells adhere and migrate on extracellular matrices thanks to their integrin receptors^23^. Many cells such as border cells in the *Drosophila* egg chamber^58^, neuronal precursors^25, 39^ or cancer cells also adhere and migrate on the plasma membrane of the adjacent cells. In this case, cell migration is mediated by cadherins which physically hold cells together^43^. Changes in expression or function of cadherins have major impacts on cell migration during neural development^59, 62, 70^ and tumour cell invasion^28, 74, 76^.

Cadherins are the homophilic ligands of *Adherens Junctions* (AJ) involved in the cohesion of solid tissues^47^. Cadherins provide anchorage between neighbouring cells thanks to their interaction with the contractile actomyosin network via catenins^46^. E-cadherin is required for epithelial cell cohesion and is recognised as a tumor suppressor^28, 73^. N-cadherin, the neuronal cadherin, although required for the cohesive interaction of neuroepithelial cells^26^, mediates weaker cell-cell adhesion and is also associated with physiological and pathological cell migration in a large range of tissues^16, 51, 65^. N-cadherin ensures weak adhesion between post-mitotic neurons and radial glial cells allowing radial neuronal migration^17, 25^. Its active endocytosis and turnover maintain proper steady-state level of N-cadherin at the cell surface allowing the effective locomotion of neurons^25^. It is also required for long distance migration of tangentially migrating interneuron precursors^39^. Moreover, N-cadherin stimulates neurite outgrowth^2, 3, 37, 42, 77^ *in vitro*. Two pathways have been involved: (i) the mechanical coupling of cadherins to actomyosin cytoskeleton, which generates the traction forces necessary to propel the growth cones^2, 19^ and (ii) the activation of FGFR-dependent biochemical signalling cascades^4, 77^.

FGFRs belong to the family of single pass transmembrane Receptor Tyrosine Kinases. Binding of their ligands, FGFs, triggers intracellular signalling cascades playing key roles during development and pathogenesis^36, 45^. Loss of expression of FGFR1 in mice disrupts the migration of epidermal cells from the primitive streak. This phenotype can be rescued by down-regulating E-cadherin-mediated intercellular adhesion^10, 11, 15, 79^ In *Drosophila*, the migration of tracheal cells requires FGFR signalling, which regulates cytoskeletal reorganisation^8, 35, 54^.

Dysfunctions of N-cadherin and FGFRs both induce pathological migrations that are most visible in cancers. N-cadherin upregulation correlates with increased motility and invasiveness of dysplastic cells in melanoma^38^, bladder^61^, prostate^31^, lung^49^ or breast cancers^48^. Mutations in FGFRs are associated to pancreatic, endometrial, bladder, prostate, lung and breast cancers^57, 75^. Literature reports on a synergistic action between N-cadherin and FGFRs in the regulation of epiblast stem cells pluripotency^67^, ovarian cells survival^71^ and osteogenic cells differentiation^13^. Overexpression of N-cadherin in mEpiSC cells prevents the downregulation of FGFR at the plasma membrane after FGF2 addition^67^. FGF and N-cadherin maintain granulosa and ovarian cells viability *in vitro* by stimulating FGFR phosphorylation^71^. The expression of a constitutively active form of FGFR increases the expression of N-cadherin reinforcing cell-cell adhesion in human osteogenic cells^13^. A functional relationship between FGFR and N-cadherin has been reported during neurite outgrowth^4, 77^ FGFR and N-cadherin co-cluster and interact at the surface of neuronal cells^4, 72^. The expression of a dominant negative FGFR inhibits neurite growth stimulated by N-cadherin^5^. In breast cancer cells, N-cadherin overexpression increases cell migration^22^. N-cadherin prevents FGFR from undergoing ligand-induced internalisation, resulting in FGFR stabilisation at the plasma membrane and sustained FGFR signalling^64^. In human pancreatic cancer xenografts, inhibition of FGFR leads to a decrease in N-cadherin expression and cell invasion^66^. Altogether, these data suggest that N-cadherin and FGFR synergise to generate signals that regulate the migratory behaviours of normal and cancer cells.

Little is known however about the combined effects of N-cadherin and FGFR activities on cell adhesion and migration. To dissect the reciprocal interplay between FGFR1 and N-cadherin, we expressed both receptors in HEK cells and analysed the consequences on N-cadherin-dependent cell adhesion and cell migration using a single cell migration model on N-cadherin coated lines. Both proteins are co-recruited and co-stabilised at cadherin-mediated cell contacts through direct interaction of their extracellular domains. As a consequence, N-cadherin-mediated cell contacts are strengthened, limiting the migration of cells on N-cadherin coated surfaces. Both the inhibition of N-cadherin-mediated migration and the stabilisation of N-cadherin at cell contacts require FGFR activity, which is itself stimulated by N-cadherin engagement. We further show that FGFR1 stabilises N-cadherin at the cell membrane by decreasing its internalisation. FGFR1 expression triggers an increase of activated Src but does not affect significantly the phosphorylated p120 catenin on tyrosine 228. However, both p120 and Src are involved in the stabilisation of N-cadherin at cell-cell contacts and in the negative regulation of N-cadherin-mediated migration induced by FGFR1. Moreover, FGFR1 stimulates the anchoring of N-cadherin to actin. Finally, we found that the migratory behaviour of cells depends on an optimum balance between FGFR-regulated N-cadherin adhesion and actin dynamics.

## Results

### FGFR1 expression inhibits N-cadherin-mediated cell migration

To study single N-cadherin-mediated cell migration and the impact on this migration on Fibroblast Growth Factor Receptor 1 (FGFR1), we developed a model in which isolated N-cadherin transfected (Ncad cells) and N-cadherin/FGFR1 double transfected HEK cells (Ncad/FGFR cells) were allowed to adhere and migrate on 10 μm width Ncad-Fc-coated stripes, in the absence of exogenously added FGF (**Fig. 1A, Video 1**). Both Ncad and Ncad/FGFR cells adhered to the surface while untransfected HEK cells did not (data not shown). Ncad cells migrated efficiently covering a total displacement up to 400 μm over 20 hours with very few inversions of direction of migration, while Ncad/FGFR cells were almost stationary (**Fig. 1B-E**). Treatment of Ncad/FGFR cells with PD173074, a FGFR kinase inhibitor, restored the migratory behaviour of Ncad/FGFR cells close to that of Ncad cells (**Fig.1A-E**), indicating that the inhibition of N-cadherin-mediated migration by FGFR1 requires the receptor kinase activity. This effect is specific to N-cadherin mediated adhesion, as we observed that the expression of FGFR1 did not impact on the migration of Ncad expressing cells on fibronectin coated surfaces (data not shown). Moreover, we verified that the blockade of FGFR activity strongly increased the migration of cells endogenously expressing N-cad and FGFR, i.e., C2C12 mouse myoblastic, U2OS human osteosarcoma and 1205Lu human metastatic melanoma (**Fig. S1**), that otherwise only moved at a similar speed as Ncad/FGFR HEK cells. Altogether these data indicate that FGFR1 strongly impairs the migration of cells on N-cadherin in a process depending on its kinase activity, but in the absence of exogenous FGF. Accordingly, further experiments were all performed in the absence of FGF.

**Figure 1:**
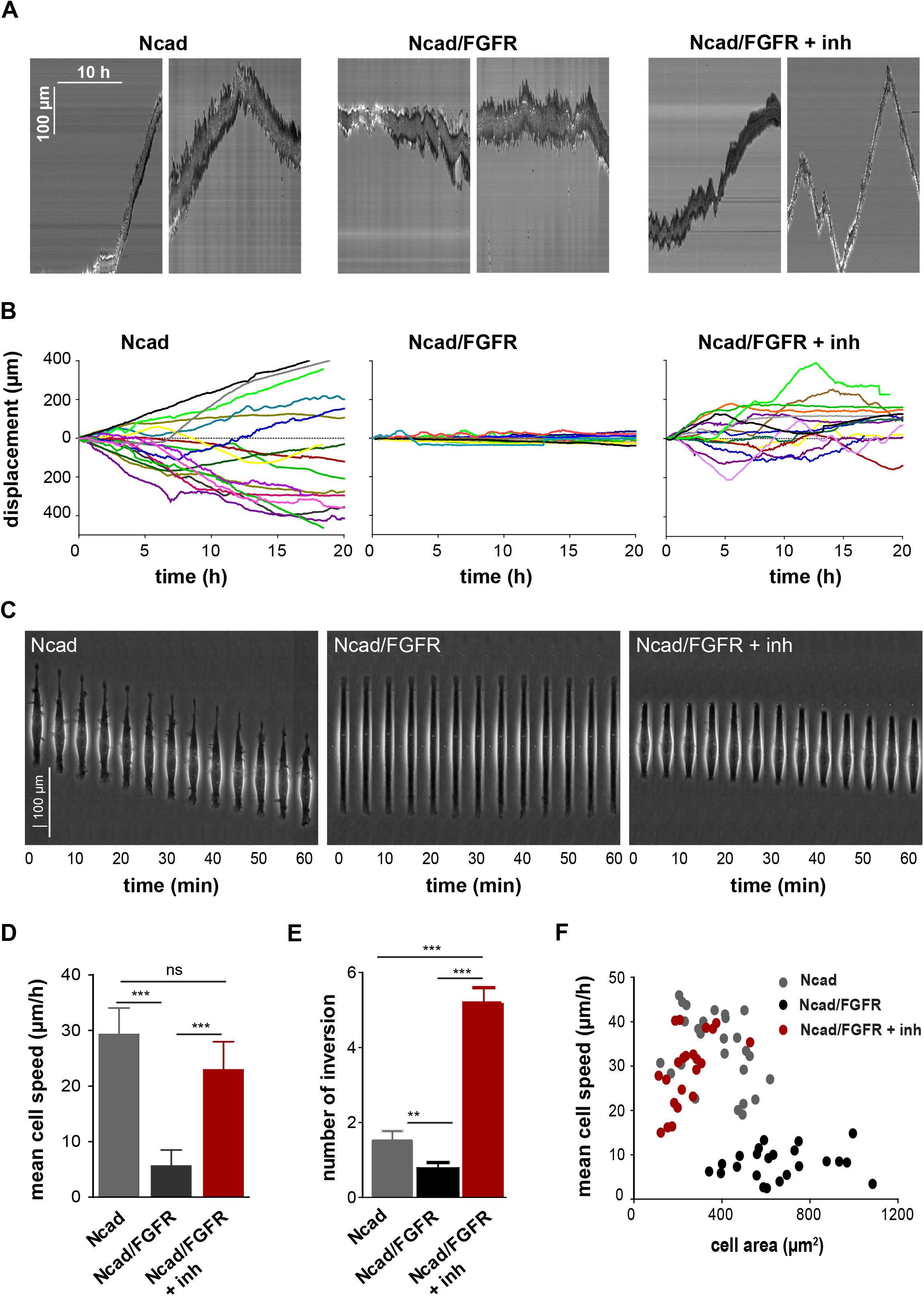
FGFR decreases the migration of N-cadherin expressing cells on N-cadherin coated lines. DsRed-Ncad (Ncad) and DsRed-Ncad/GFP-FGFR (Ncad/FGFR) expressing HEK cells were seeded at low density on 10 μm-width Ncad-Fc coated lines in the absence or in the presence of FGFR inhibitor (Ncad/FGFR+inh) and imaged in phase contrast every 6 minutes during 20 hours (see Video 1). **(A)** Representative kymographs of the displacement over 10 hours of two cells for each condition. **(B)** The trajectories of single cells Ncad (n = 26), Ncad/FGFR (n = 22) and Ncad/FGFR+inh (n = 25) were manual tracked and the cell displacement plotted over time. **(C)** Representative kymographs of 1-hour long cell displacements imaged at higher magnification (see Video 1). **(D, E)** Histograms representing the mean cell body speed and the frequency of inversion in migration direction, respectively, for Ncad, Ncad/FGFR and Ncad/FGFR+inh cells (** p ≤ 0.01, *** p ≤ 0.0001, ANOVA multi-comparison test, Newman-Keuls post-test). **(F)** Plots of the mean cell body speed as a function of cell surface area for the three cell populations.

Ncad/FGFR cells were more spread than Ncad cells, a trend that was reverted in the presence of the FGFR inhibitor (**Fig. 1C, Video 1**). The mean spreading area of Ncad/FGFR cells of 764.7 ± 24.9 μm^2^ was reduced to 562.6 ± 10.6 μm^2^ in the presence of the inhibitor, which is close to the values measured for Ncad cells (302.8 ± 13.4 μm^2^). Plotting the mean cell speed as a function of cell area confirmed an inverse correlation between these two parameters: the more the cells spread on N-cadherin, the slower they migrate (**Fig. 1F**). Ncad/FGFR cells displayed an extensive spreading and a reduced migration speed. Thus, the reduced migration of FGFR1 expressing cells could result from a strengthening of cadherin-mediated adhesion on the Ncad-Fc coated lines.

### N-cadherin and FGFR1 are co-stabilised at cell-cell contacts

We hypothesised that FGFR1 increases N-cadherin-mediated cell-cell adhesion by affecting the dynamics of junctional N-cadherin. To test this hypothesis, we performed dual wavelength FRAP (Fluorescence Recovery After Photobleaching) experiments at cell-cell contacts of cells expressing DsRed-Ncad or GFP-FGFR1 (alone) or both, in the absence of added FGF (**Fig. 2A, B**). Expressed alone, N-cadherin displayed a mobile fraction at the cellcell contacts of 61.3 ± 2.7 %. Coexpression of FGFR1 decreased this value to 38.2 ± 3.4 % while treatment with the FGFR inhibitor restored N-cadherin mobile fraction level (62.3 ± 2.3 %) close to that of Ncad cells. Expression of N-cadherin also significantly decreased the mobile fraction of junctional FGFR1 (**Fig. 2C**). Similar experiments were performed using mCherry-tagged E-cadherin (mCherry-Ecad) instead of DsRed-Ncad, revealing that the E-cadherin expression did not affect the dynamics of FGFR1 at cell-cell contacts (**Fig. 2C**). FGFR1 expression did not modify the mobile fraction of E-cadherin at cell-cell contacts (**Fig. 2D**). Thus, FGFR1 and N-cadherin specifically co-stabilise each other at N-cadherin-mediated contacts. This co-stabilisation may lead to the strengthening of N-cadherin adhesion, which may explain the increased spreading of cells on Ncad-coated lines upon FGFR1 expression.

**Figure 2:**
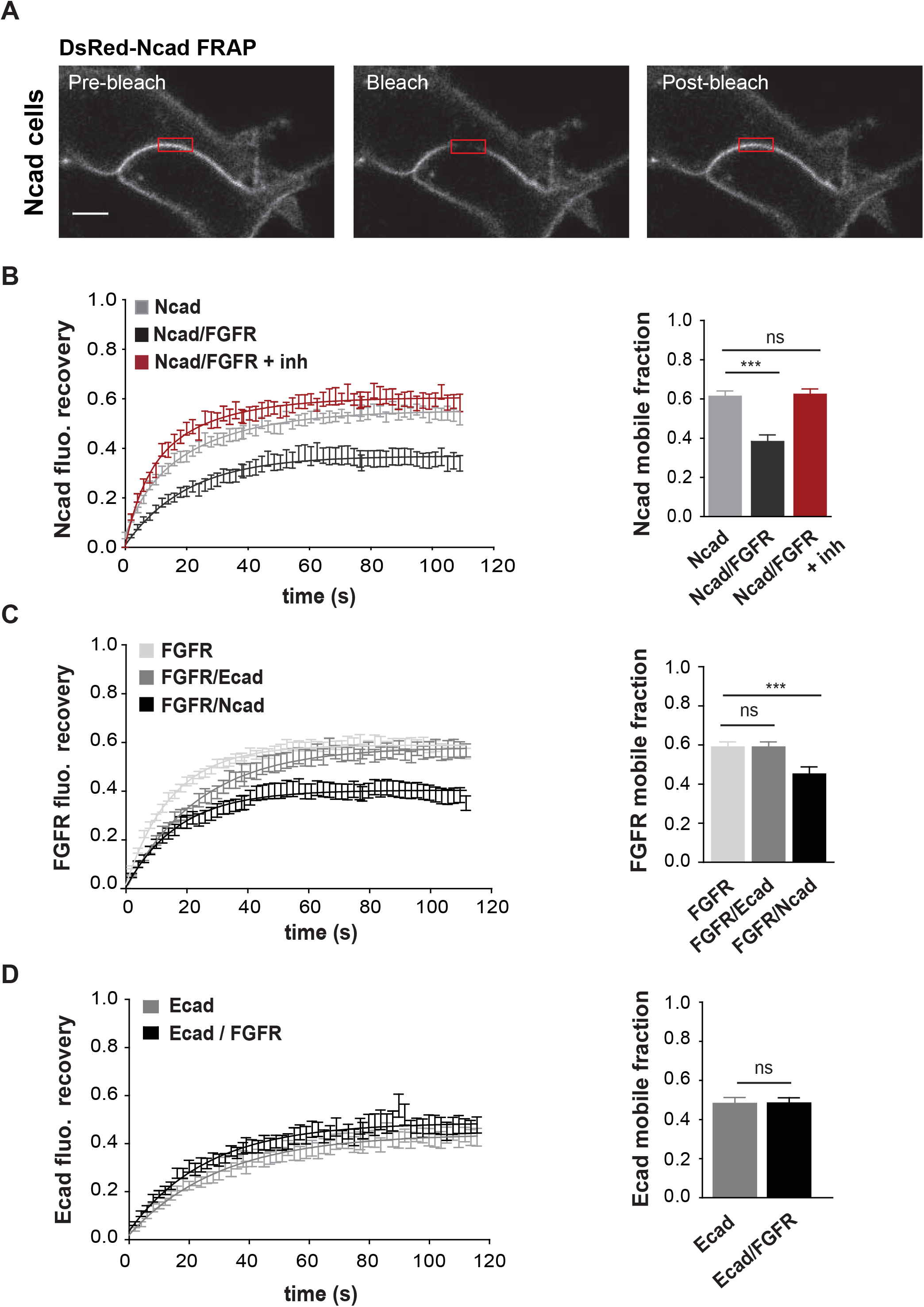
FGFR and N-cadherin co-stabilise each other at cell-cell contacts. **(A)** Representative images of FRAP experiments performed at the cell-cell contacts of DsRed-Ncad HEK cells: Fluorescent signal before (Pre-bleach), immediately after bleaching (Bleach) and 110 sec after the bleach (Post-bleach). Red rectangles represent the bleached region at cell-cell contacts. Scale bar: 40 μm. **(B)** Left: normalised DsRed-Ncad fluorescence recovery curves for Ncad, Ncad/FGFR and Ncad/FGFR+inh cells, respectively. n = 28 independent experiments. Right: mean Ncad mobile fraction ± SEM, *** p ≤ 0.001; ns: nonsignificant, ANOVA multiple comparison test, n = 28 from 3 independent experiments. **(C)** Left: normalised fluorescence recovery for GFP-FGFR in FGFR, FGFR/Ecad and FGFR/ Ncad cells, respectively. n = 15. Right: mean FGFR mobile fraction ± SEM), *** p≤ 0.0001; ns: non-significant, ANOVA multi-comparison test, n = 15 from 3 independent experiments. **(D)** Left: normalised mCherry-Ecad fluorescence recovery in mCherry-Ecad (grey) and mCherry-Ecad/FGFR (black) cells, respectively. n = 18 from 3 independent experiments, Right: mean mCherry-Ecad mobile fraction ± SEM, ns: non-significant, student t-test, n = 18.

### FGFR1 stimulates junctional N-cadherin accumulation and strengthens cell-cell contacts

To confirm the strengthening effect of FGFR1 on N-cadherin-mediated adhesion, we quantified the accumulation of N-cadherin at cell-cell contacts in cell monolayers. Cell-cell contacts accumulated more N-cadherin and were straighter for Ncad/FGFR than for Ncad cells (**Fig. S2A**). FGFR inhibitor treatment inhibited the effect of FGFR1 on both the straightness and junctional accumulation of N-cadherin, indicating that the kinase activity of the receptor is required for the strengthening of N-cadherin-mediated cell contacts. Accordingly, analysing N-cad distribution at cell-cell contacts in cell doublets grown on fibronectin-coated lines, as well as in suspended cell doublets, revealed an increased accumulation of junctional N-cadherin in the presence of FGFR1 (**Fig. S2B, S3**).

To determine the impact of this junctional N-cadherin stabilisation on cell-cell contact stability, we followed by live imaging the disassembly of cell-cell contacts upon chelation of Ca^2+^ ions in cell monolayers (**Fig. 3A**). Quantitative analysis of cell-cell contact life-time following Ca^2+^ depletion indicated that Ncad cells were dissociated in 2 minutes while it took almost 6 minutes to dissociate Ncad/FGFR cells. This cell-cell contact stabilisation was prevented by FGFR activity inhibition. Notice that the inhibitor had no effect on the dissociation of intercellular contacts of Ncad cells. To test more directly the effect of FGFR1 on the strength of N-cadherin-mediated adhesion, we probed the response to force developed at contacts between Ncad-Fc-coated magnetic beads and Ncad or Ncad/FGFR cells. Beads were left to interact with the cell surface for 30 minutes, before being probed for displacement under force by approaching a magnetic rod. After calibration, one can estimate the actual forces at which the N-cadherin-mediated adhesions between the bead and the plasma membrane were disrupted (**Fig. 3B, Video 2**). Significantly fewer beads were displaced or detached from the cell surface of Ncad/FGFR cells indicating that the binding strength was higher compared to Ncad cells. The inhibition of the FGFR activity restored bead detachment/displacement level close to the one observed for Ncad cells (**Fig. 3C**). For the population of beads that were detached under force, the mean breaking distance was of 28.5 ± 0.9 μm for Ncad cells and 14.3 ± 0.6 μm for Ncad/FGFR cells, respectively. Inhibition of FGFR increased the breaking distance to 21.4 ± 0.9 μm in Ncad/FGFR cells, corresponding to rupture forces of 5.9 ± 0.1 nN, 7.3 ± 0.1 nN and 6.5 ± 0.1nN, respectively. Thus, FGFR1 and its kinase activity increase the mechanical resistance of N-cadherin adhesion.

**Figure 3:**
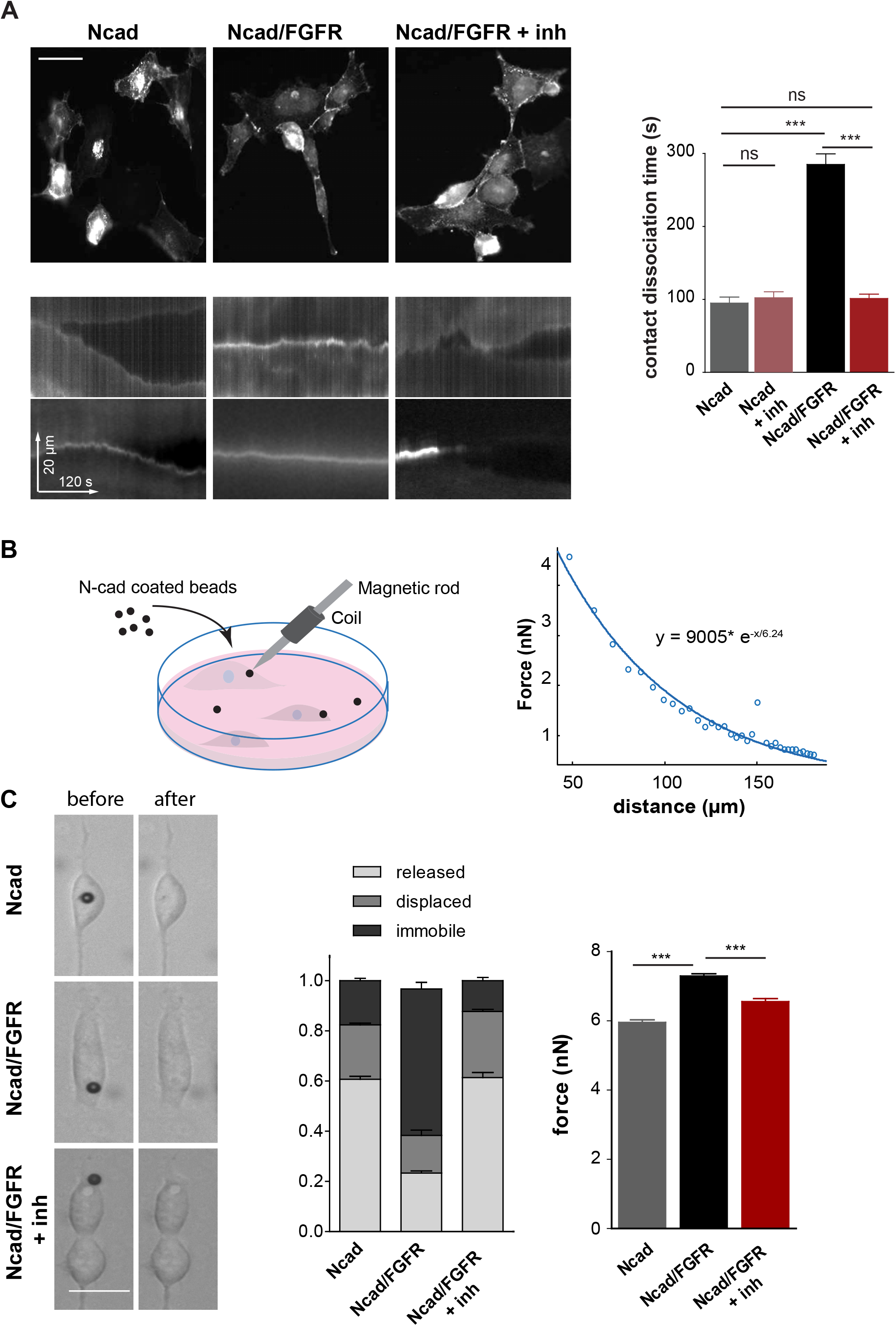
FGFR strengthens N-cadherin-mediated cell-cell contacts and reinforces N-cadherin anchoring to the cell cortex. **(A)** Ncad, Ncad/FGFR, Ncad/FGFR + inh cells cultured at confluence over 1 mm^2^ square fibronectin-coated-patterns were treated with EGTA then imaged for DsRed-Ncad every 30 seconds during 15 min. **(top left)** Low magnification images taken after 5 min of EGTA treatment. Scale bar = 40 μm. **(buttom left)** Examples of kymographs of the DsRed-Ncad signal along a line perpendicular to the cell-cell contact starting from EGTA addition (t_0_) for Ncad, Ncad/FGFR and Ncad/FGFR+inh cells. **(right)** Contact dissociation time upon EGTA addition as determined from the kymographs for the three conditions, plus Ncad+inh cells. ** p≤0.01; *** p≤0.001, ANOVA multi comparison test, Newman-Keuls post-test, n = 60 contacts. **(B)** Left: magnetic tweezers experimental set up used to evaluate the anchorage and rupture force of N-cadherin mediated bead-cell contacts. Right: Calibration curve of the magnetic tweezers determined as described in material and methods. The force is exponentially inversely correlated with the bead-magnetic needle distance. **(C, left)** Representative images of Ncad-coated beads before and after tweezer-induced detachment from the cell membrane. **(middle)** Distribution of the responses of Ncad beads to the magnetic field in three classes (release, displacement, and immobility) for Ncad (n = 60), Ncad/FGFR (n = 65) and Ncad/FGFR+inh (n = 50) cells. **(right)** Bead-cell contact disruption forces calculated from the Stoke equation for Ncad, Ncad/FGFR and Ncad/FGFR + inh HEK cells (*** p ≤0.001, ANOVA multi comparison test, Newman-Keuls post-test).

### N-cadherin and FGFR1 interaction promotes FGFR1 activation

We described so far the effects of FGFR1 on N-cadherin-mediated adhesion and migration, both requiring the kinase activity of the receptor although no exogenous FGF ligand was added. Furthermore, FGFR1 and N-cadherin co-localised and co-stabilised at the cadherin-mediated cell contacts. Therefore, we hypothesised that the increased residence of FGFR at cell-cell contacts induced by N-cadherin-mediated adhesion could induce an activation of the receptor relying on direct interaction of these two proteins, as previously reported in neuronal cells^4^. To confirm this hypothesis, the level of binding of Ncad-Fc to immobilised FGFR1 extracellular domain was measured using an optical biosensor. Results showed a direct interaction between N-cadherin and FGFR1 ectodomain with an affinity, calculated from the kinetic parameters of the interaction, of 106 ± 25 nM) (**Fig. 4A and Suppl. Table I**). This interaction was confirmed by co-immunoprecipitation of GFP-FGFR1 from protein extracts of HEK cells co-expressing the two proteins (**Fig. 4B**). The coprecipitation was strongly reduced when FGFR kinase activity was inhibited.

**Figure 4:**
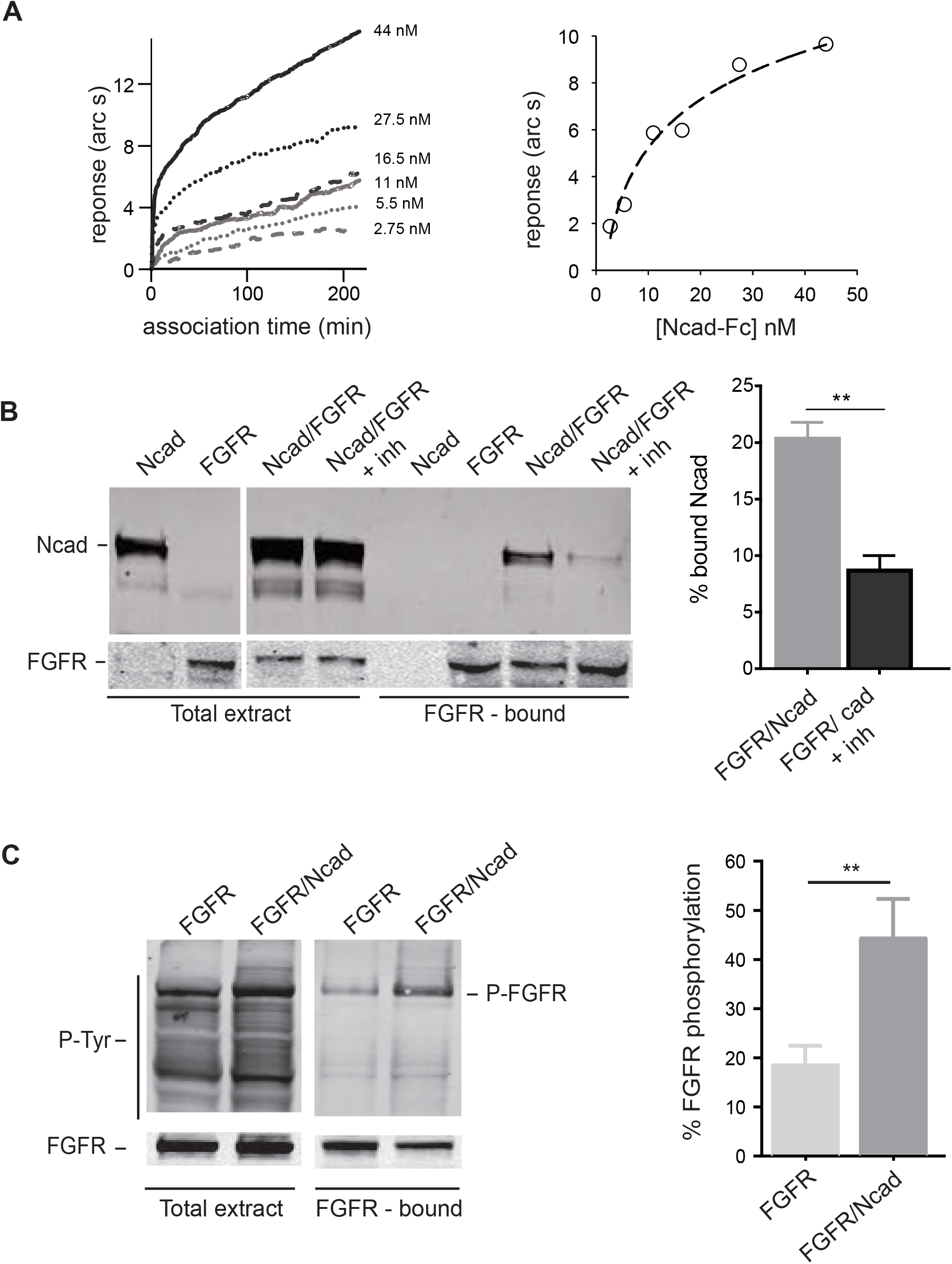
N-cadherin and FGFR associate leading to increased activation of FGFR. **(A)** Binding of Ncad-Fc to FGFR1 extracellular domain. Kinetics of Ncad-Fc to immobilised FGFR1 extracellular domain was measured as described under “Materials and Methods.” Left panel: Ncad-Fc at different concentrations was added to a FGFR1-derivatised cuvette and the association reaction was followed for 200 s. Data were collected three times a second. The concentration of Ncad-Fc is indicated. Data shown are the result of one representative experiment out of three. Right panel: relationship between the extent of binding (response in arc s) of the association reactions shown in left and Ncad-Fc concentration. All results are summarised in supplementary table 1. **(B)** GFP-FGFR was immunoprecipitated with anti-GFP bead from protein extracts of Ncad, FGFR and Ncad/FGFR cells. Immunoprecipitates, together with total protein extracts, were then analysed by Western blot using anti-Ncad and anti-GFP (FGFR) antibodies. The histogram shows the ratio of N-cadherin bound to GFP-FGFR on N-cadherin in total extract, determined from the quantification of 3 independent immunoblots then converted to percentage. ** p≤0.01, Student’s t test, n = 3. **(C)** To detect FGFR phosphorylation GFP-FGFR immuno-precipitates were immunoblotted with anti-P-Tyr and anti-GFP (FGFR) antibodies. The histogram shows the ratio of P-Tyr on GFP-FGFR signals as a quantification of the degree of phosphorylation of FGFR in the different extracts. ** p ≤ 0.01, Student *t* test, n = 3.

Then, we tested whether N-cadherin could induce FGFR activation in the absence of exogenous FGF. Our results revealed that FGFR1 phosphorylation was significantly increased in Ncad/FGFR cells compared to cells only expressing FGFR1 (**Fig. 4C**). To further provide evidence that FGFR1 was activated by N-cadherin-mediated adhesion, we probed the activation of Erk1/2, a well-known downstream effector of FGFR signalling, following Ca^2+^ switch in C2C12 cells that express endogenous N-cadherin^18^ and FGFRs^32^ (**Fig. S4**). Addition of Ca^2+^ for 10 minutes to Ca^2+^-depleted cells significantly increased Erk1/2 phosphorylation in the absence, but not in the presence of the FGFR inhibitor, strongly suggesting that N-cadherin engagement triggers the activation of the FGFR1. Altogether our results suggest a two-way communication between FGFR1 and N-cadherin resulting from their direct interaction. The stabilisation of FGFR1 by N-cadherin at cell-cell contacts allows its activation. The activation of FGFR1 could in turn increase junctional N-cadherin stabilisation, responsible for the observed strengthening of N-cadherin-mediated cell adhesion and reduction of N-cadherin-dependent cell migration.

### FGFR1 stabilises N-cadherin at the plasma membrane through downregulation of endocytosis

To determine whether FGFR1 expression also increases N-cadherin prevalence at the plasma membrane, we performed cell surface biotinylation on Ncad and Ncad/FGFR HEK cells. The fraction of cell surface exposed biotin-labelled N-cadherin was significantly higher in Ncad/FGFR cells than in Ncad cells. It was strongly decreased in Ncad/FGFR cells following treatment with the FGFR inhibitor (**Fig. 5A**). Thus, FGFR1 favours the accumulation of N-cadherin at the plasma membrane in a process depending on its kinase activity. A first hint on the way FGFR1 may regulate N-cadherin availability at the cell surface was given by imaging DsRed-Ncad and analysing its distribution in Ncad or Ncad/FGFR cells thanks to flow cytometry imaging (**Fig. 5B**). Accordingly, when imaging DsRed-Ncad in cells migrating on fibronectin-coated lines (**Video 3**), we observed N-cadherin vesicles trafficking from the leading edge to the rear of the cells. These vesicles were more prominent in Ncad than in Ncad/FGFR cells, suggesting that the trafficking of N-cadherin was reduced in the latter (**Fig. 5C**). We thus questioned the role of endocytosis in the regulation of cell surface N-cadherin by FGFR1 by quantifying N-cadherin endocytosis following the internalisation of biotinylated cell surface proteins (**Fig. 5D, Fig. S5**). The N-cadherin endocytic pool was significant reduced in Ncad/FGFR cells compared to Ncad cells. This effect was significantly reduced in the presence of FGFR inhibitor, although the inhibition was far from complete (**Fig. S5**). Moreover, treatment of Ncad and Ncad/FGFR cells with hydroxyl-dynasore, an endocytosis inhibitor decreased the fraction of endocytosed N-cadherin in Ncad cells to levels measured for Ncad/FGFR cells (**Fig. 5D**). These data support the notion that FGFR1 upregulates N-cadherin prevalence at the plasma membrane by inhibiting its endocytosis, a process that could contribute to the reinforcement of N-cadherin-mediated cell contacts.

**Figure 5:**
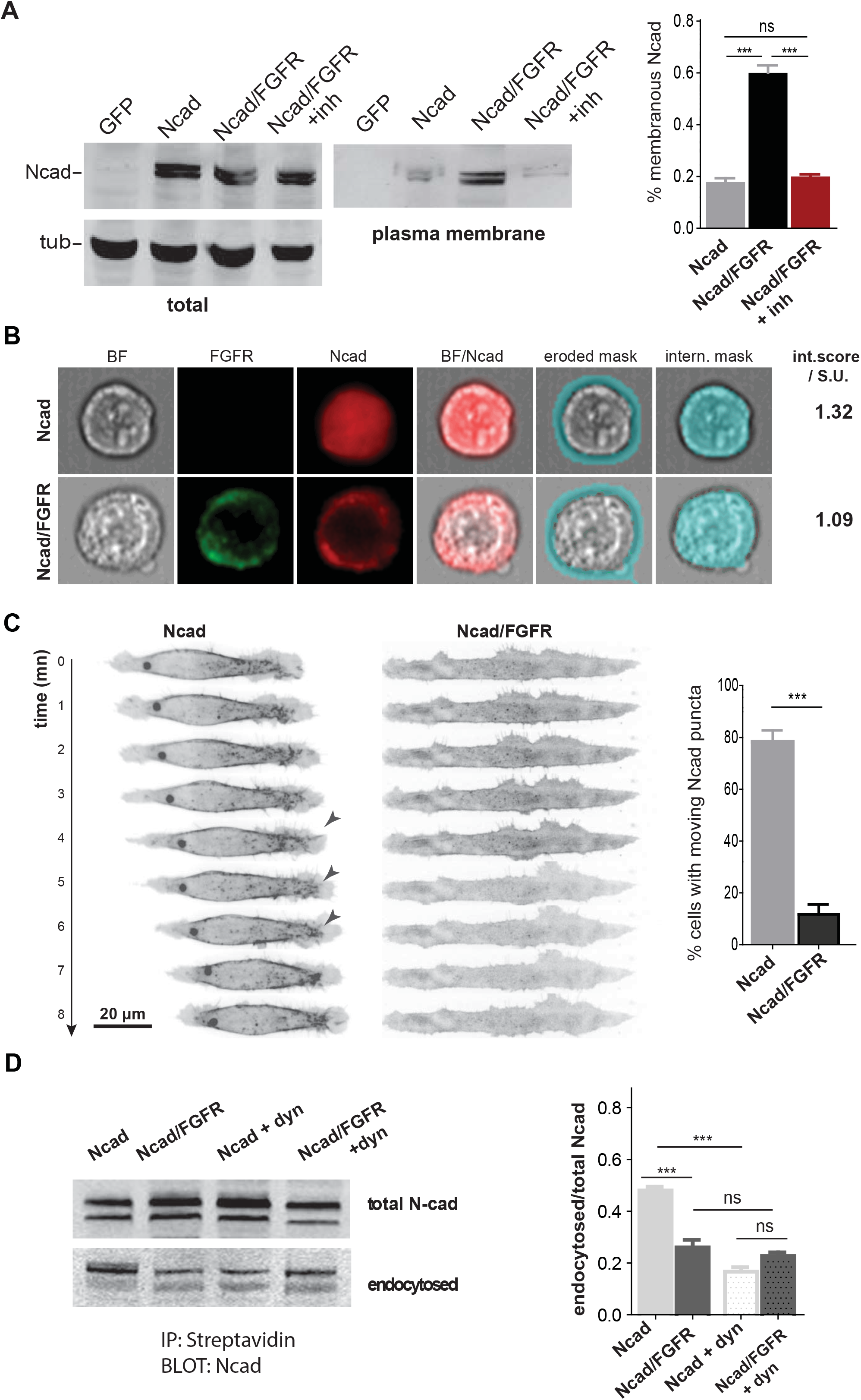
FGFR increases N-cadherin cell surface accumulation by reducing its endocytosis. **(A)** Analysis of cell surface expression of N-cadherin. After surface biotinylation at cold and removal of unfixed biotin, Ncad, Ncad/FGFR and Ncad/FGFR+inh cells were immediately lysed and protein extracts subjected to precipitation by streptavidin beads. GFP transfected HEK cells were used as control. Total extracts and streptavidin bound fractions (plasma membrane exposed fractions) were then immunoblotted with anti-N-cadherin antibodies. The histogram shows the quantification of N-cadherin exposed at the plasma membrane over total N-cadherin content for the three conditions. *** p ≤ 0.001, ns: non-significant, ANOVA multi comparison test, Newman-Keuls post-test, n = 4. **(B)** Analysis of Ncad internal pool by flow cytometry imaging. Ncad and Ncad/FGFR cells were non-enzymatically detached, then processed for flow cytometry imaging in bright field, and for dsRed-Ncad and GFP-FGFR fluorescence imaging. Masks were defined on bright field images to separate cell membrane and internal cell areas on each cell. Applied to the fluorescence images they allowed to extract an internalisation score as described in Materials & Methods. FGFR reduces the internalisation score of N-cadherin molecule by 17% (1.09 U.I versus 1.32 U.I). Experiences were repeated 4 times, over populations of 150.000 cells for each condition in each experiment. **(C)** Ncad and Ncad/FGFR cells were seeded on Ncad-coated stripes of 10 μm, then after4 hours, preparations were imaged at 63X for Ds-Red Ncad. The panels show the maximum projection of 1μm thick confocal sections encompassing the whole cell thickness. Arrow-heads show N-cadherin puncta trafficking from the leading edge to the rear of Ncad expressing cells. The histogram shows the quantification of the percentage of cells with such puncta. **p ≤ 0.01; non parametrical t test; n = 15, n = 20 cells for Ncad and Ncad/FGFR cells, respectively. **(D)** Analysis of N-cad endocytic fraction following cell surface biotinylation. Freshly biotinylated Ncad and Ncad/FGFR cells were switched to 37°C for 40 minutes to allow endocytosis to resume in the presence or in the absence of dynasore, then subject to a reducing wash in order to remove remaining medium exposed biotin. Left: cells were lysed and protein extracts subjected to precipitation by streptavidin beads, then anti-N-cadherin bound (total) and streptavidin bound fractions (endocytosed) were immunoblotted with anti-N-cadherin antibodies. Right: The histogram shows the ratio of endocytosed over total Ncad in each extract. *** p ≤ 0.001; ns: non-significant, ANOVA multiple comparison test, n = 3 experiments.

### The effects of FGFR1 on N-cadherin-mediated adhesion and migration involves p120

p120 has been reported to stabilise cadherins at cell-cell contacts by regulating their trafficking^24^ either to the plasma membrane^7^ or from the plasma membrane to endocytic compartments^12, 78^. In particular, it would do so by binding and masking an endocytic signal conserved in classical cadherins^50^. We thus asked whether the interaction of N-cadherin with p120 was involved in the regulation of N-cadherin endocytosis by FGFR1. First, we measured the intensity of GFP-p120 fluorescence along Ncad or Ncad/FGFR cells in contact (doublets). The results showed an increased recruitment of GFP-p120 at cell-cell contacts in Ncad/FGFR cells compared to Ncad cells (**Fig. 6A**). Then, to test the implication of p120, we coexpressed FP tagged FGFR1 and the NcadAAA mutant. The AAA mutation at position 764 in E-cadherin^68^ and N-cadherin^69^ was described to impair their binding to p120. FRAP experiments on Ncad/FGFR and NcadAAA/FGFR cells revealed that the mobile fraction of the mutated N-cadherin was significantly higher than that of the wild type molecule (**Fig. 6B**), suggesting that the binding of p120 to N-cadherin was involved in the stabilisation of N-cadherin induced by FGFR1. To test whether the ability of N-cadherin to bind p120 also affects N-cadherin-mediated cell migration, we compared the migration of Ncad/FGFR and NcadAAA/FGFR cells (**Fig. 6C, D and Video 4**). NcadAAA/FGFR cells displayed reduced spreading areas and increased migration speeds compared to Ncad/FGFR cells (**Fig. 6C**). They migrated at a speed very similar to the one of Ncad cells (see **Fig 1D**). Interestingly, the mutation drastically reduced the propensity of N-cadherin to interact with FGFR1 (**Fig. S6**). Thus, preventing the binding of N-cadherin to p120 strongly inhibits the junctional N-cadherin stabilisation and the single cell migration inhibition induced by FGFR1.

**Figure 6:**
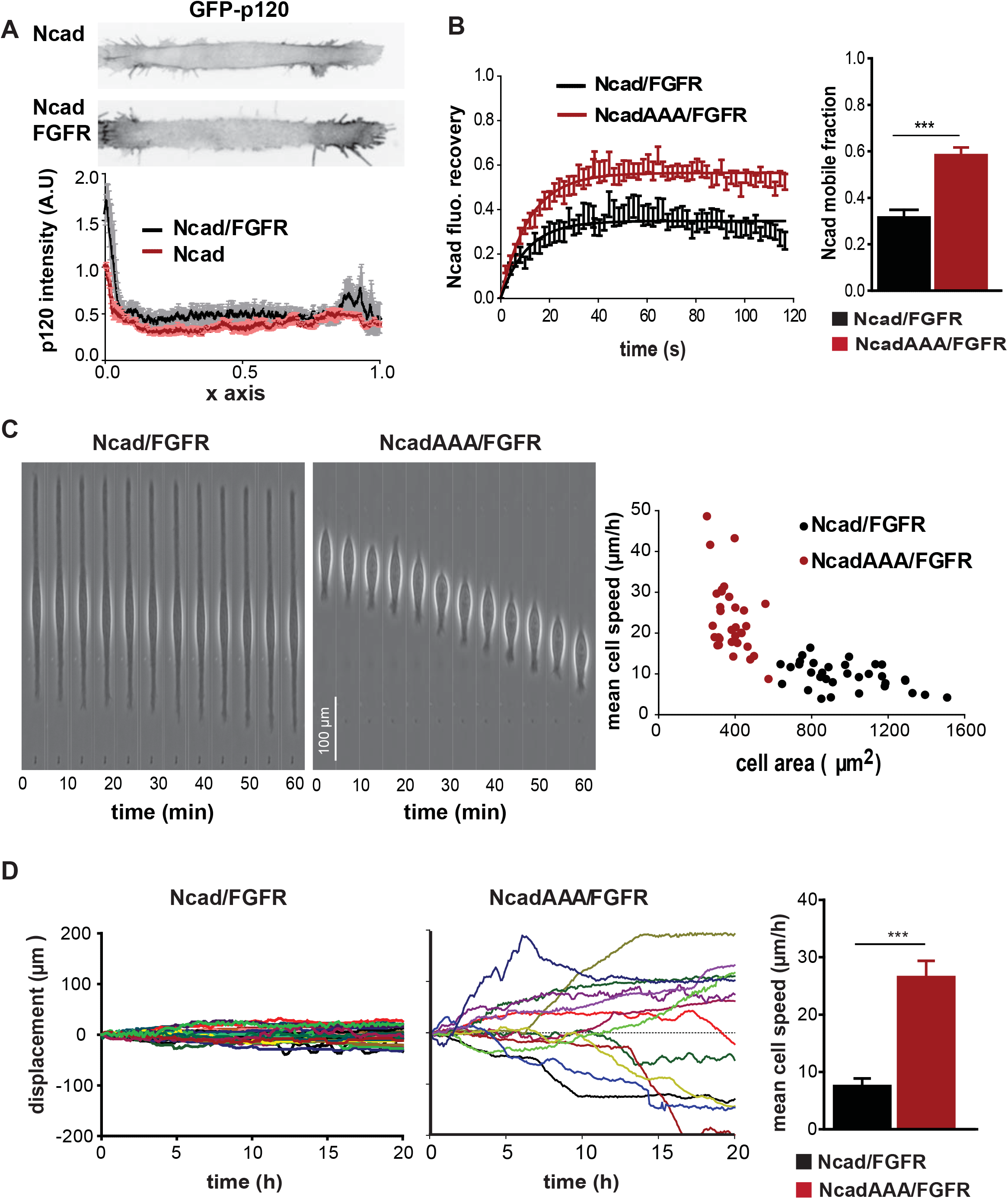
p120 is involved in the stabilisation of N-cadherin at cell-cell contacts and the decreased migration induced by FGFR expression. **(A)** Ncad and Ncad/Flag-FGFR cells were transfected with GFP-p120 and seeded on fibronectin coated lines. Mean p120 intensities along the cell length with 0 as junctional end and 1 as free end of the cell was calculated on 25 cell doublets. p120 junctional accumulation was higher in Ncad/FGFR cell doublets than in Ncad doublets. **(B)** FRAP experiments were performed on cells expressing GFP-FGFR1 and either DsRed-Ncad or mCherry-NcadAAA. Curves show Ncad and NcadAAA normalised fluorescence recoveries over time for Ncad/FGFR (black) and NcadAAA-FGFR (red) cells (mobile fraction 0.29 ± 0.1 and 0.50 ± 0.1, respectively (n = 25), (***, p ≤ 0.0001, Student t test). **(C)** Ncad/FGFR and NcadAAA/FGFR cells were seeded on Ncad-Fc coated stripes and imaged every 6 minutes during 20 hours. Left: examples of Ncad/FGFR and NcadAAA/FGFR individual cell displacements over 1 hour. Right: histograms representing the mean cell speeds as a function of cell areas. **(D)** Plots show the displacement in function of time for Ncad/FGFR (left), NcadAAA/FGFR (right) cells with respectively n = 30, n = 40 cells. Histograms show the mean speed of Ncad/FGFR (black), NcadAAA/FGFR (red) cells (****, p ≤ 0.0001, ANOVA multi-comparison test).

Next, we analysed the expression levels of p120 in Ncad, FGFR and Ncad/FGFR cells (**Fig. S7A**). However, the cellular levels of p120 were not significantly affected by the expression of FGFR1. The N-terminal phosphorylation domain of p120, containing tyrosine residues phosphorylated by Src family kinases, has been reported to regulate negatively N-cadherin stability at the plasma membrane^24, 34, 40^. We thus analysed the phosphorylation on Y228 of p120 (**Fig. S7B**). However, no significant effect of FGFR1 on the phosphorylation of p120 on this site was observed. Thus, although p120 has been reported as a substrate of Src^27, 40^, itself a well-known downstream target of FGFR1^60, 80^, the involvement of p120 in regulating N-cadherin trafficking upon FGFR1 may not rely on a Src-dependent tyrosine phosphorylation pathway. Accordingly, Src inhibition by PP2 had no effect on the level of p120 phosphorylation on Y228 in Ncad/FGFR cells (**Fig. S7B**).

### The effects of FGFR1 on N-cadherin-mediated adhesion and migration involve Src family kinases

We then investigated the levels of Src activation in N-cadherin immunocomplexes in Ncad and Ncad/FGFR cells (**Fig. 7A**). FGFR1 expression led to an increase of the phosphorylation in the Src catalytic domain, while FGFR inhibition prevented this increase, suggesting that FGFR1 kinase activity is responsible for the increase in activated Src associated to N-cadherin. In order to determine the involvement of Src in the stabilisation of N-cadherin-mediated adhesion induced by FGFR1 expression, we analysed the effect of Src inhibition on the mobility of junctional N-cadherin. FRAP experiments revealed that the inhibition of Src by PP2 restored high levels of mobile junctional N-cadherin in Ncad/FGFR cells, comparable to those found in cells that do not express the receptor (**Fig. 7B**). When Ncad/FGFR cells were submitted to the single cell migration assay on N-cadherin in the presence of PP2 they displayed a strong stimulation of their migration properties with a mean speed of migration comparable to the one of Ncad cells (**Fig. 7C and Video 5**). Thus, Src inhibition counteracts both the stabilisation of N-cadherin cell-cell contacts and the inhibition of N-cadherin-mediated migration induced by FGFR1 expression. These data suggest that the activation of Src by FGFR1 in N-cadherin complexes may regulate the stability of junctional cadherin and the migratory response of N-cadherin expressing cells.

**Figure 7:**
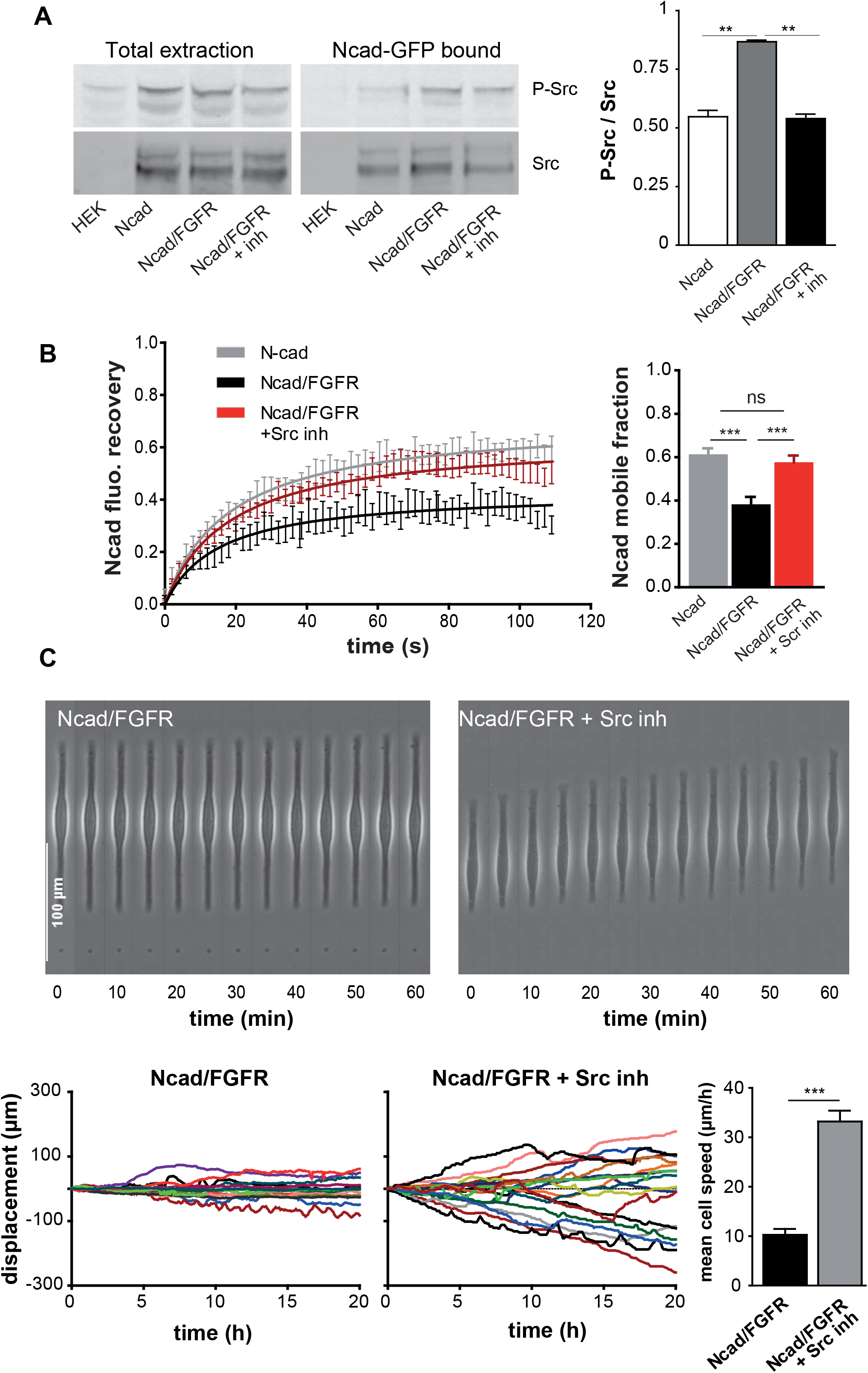
Src activity is involved in the stabilisation of N-cadherin at cell-cell contacts and the decreased migration induced by FGFR expression. **(A)** Western blot detection of N-cad, Src and phosphorylated Src (P-Src) in the Ncad immunoprecipitates. Histogram shows the ratio of phosphorylated Src calculated as the ratio of P-Src band intensity on Src band intensity. **(B)** FRAP experiments were performed on DsRd-N-cadherin at cell-cell contacts of Ncad and Ncad/FGFR in the absence or in the presence of Src inhibitor. Curves and histograms show Ncad and NcadAAA normalised fluorescence recoveries over time and extracted mobile fractions ± SEM, *** p ≤ 0.001; ns: non-significant, ANOVA multiple comparison test, n = 18). **(C)** Migration of Ncad/FGFR and Ncad/FGFR cells treated with the Src inhibitor on Ncad-Fc coated lines. Graph shows the cumulative cell displacements in function of time and histogram the mean cell migration speeds. (** p ≤ 0,01, *** p ≤ 0.0001, ANOVA multi-comparison test, Newman-Keuls post-test).

### FGFR1 stiffens the anchoring of N-cadherin to actin network

The anchoring of cadherins to actomyosin has been reported as a major mechano-signalling leading to cell-cell contact reinforcement and neuronal cell migration^19, 46^. To evidence the effect of FGFR activity on the mechanical link between N-cadherin and actin, we analysed the retrograde flow of actin in the lamellipodia of LifeAct-GFP expressing C2C12 myogenic cells spread on N-cadherin as a proxy of the coupling of cadherin to the treadmilling actin^56, 63^. The speed of F-actin rearward flow was increased by 40% in cells treated with the FGFR inhibitor compared with cells treated by the vehicle alone (**Fig. 8A; Video 6**), indicating that FGFR activity stimulates the coupling of N-cadherin to the actin cytoskeleton.

**Figure 8:**
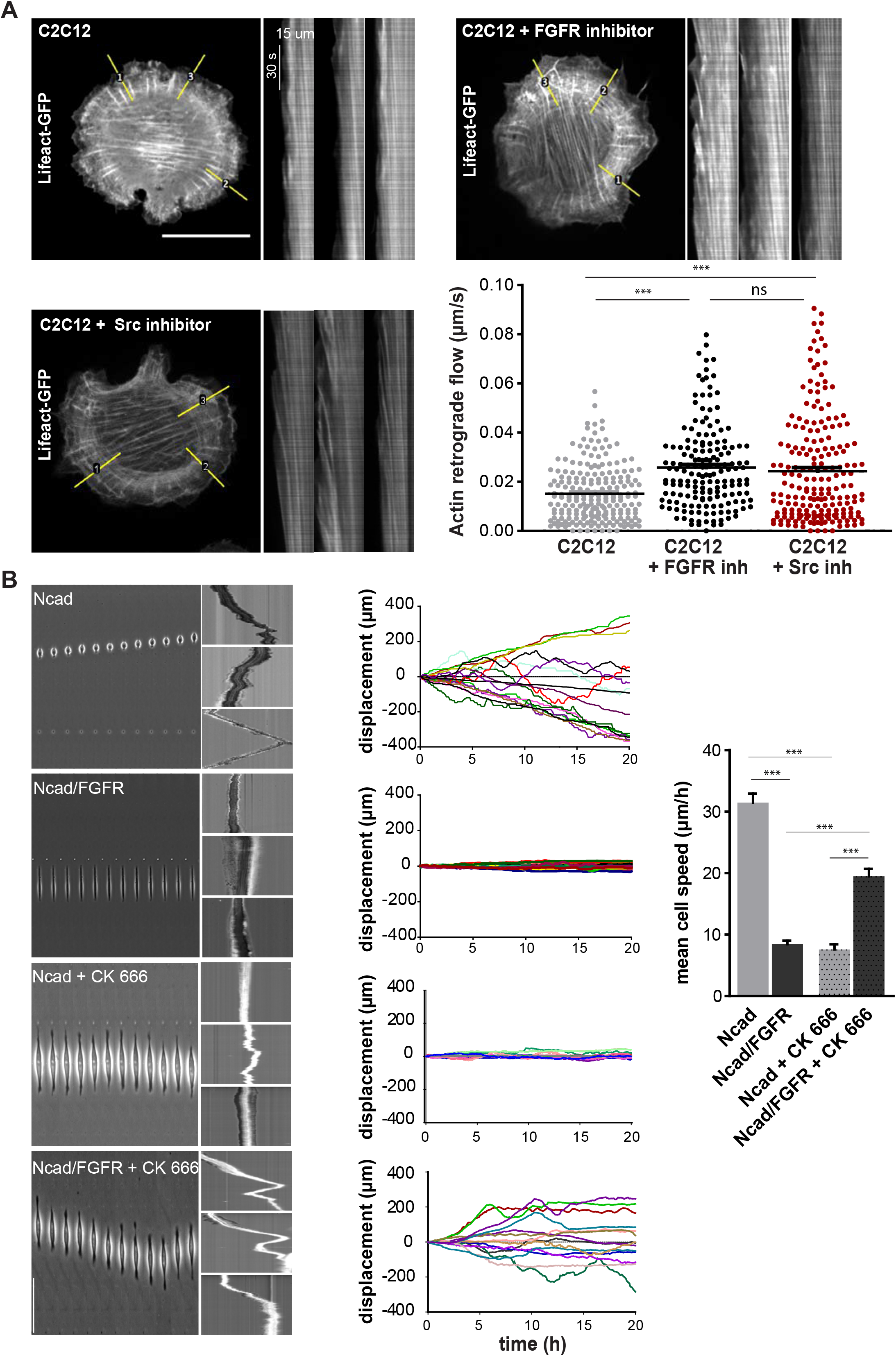
FGFR promotes N-cadherin-F-actin functional mechanocoupling. **(A)** LifeAct-GFP expressing C2C12 cells were seeded on Ncad-Fc coated surfaces for 2 hours, than treated with or without FGFR inhibitor for 1 hour, and then imaged for 5 minutes at a frequency of two images per second. Left: still images of the LifeAct-GFP signal, scale bar = 20 μm. Inserts on the right represent examples of kymograph constructed along the two pixel-wide yellow lines (1-3), Right: the actin retrograde flow was quantified by kymograph analysis. Right: the histogram shows the mean actin retrograde flow speed for C2C12 (n = 140 kymographs from 24 cells) and C2C12 + inh (n = 156 kymographs from 25 cells) cells, (**** p ≤ 0.0002, Student *t* test). **(B)** Migration on Ncad-Fc coated lines of Ncad and Ncad/FGFR cells treated or not with the Arp2/3 inhibitor. Left: kymographs of the displacement over 10 hours of three cells for each condition. Middle: cumulative displacements of cells in function of time (20 hours) for Ncad (n = 25), Ncad/FGFR (n = 25), Ncad + CK666 (n = 26), Ncad/ FGFR + CK666 (n = 22) conditions. Right: histograms representing the mean cell speed for each condition (** p ≤ 0,01, *** p ≤ 0.0001, ANOVA multi-comparison test, Newman-Keuls post-test).

To test whether this mechanocoupling modulation was instrumental in regulating N-cadherin-mediated cell migration, Ncad and Ncad/FGFR HEK cells were treated with the Arp2/3 inhibitor CK 666 and analysed for their migration on Ncad-coated lines (**Fig. 8B; Video 7**). While the inhibition of branched actin polymerisation almost fully abrogated the migration of Ncad cells, it significantly increased the migration of Ncad/FGFR cells on N-cadherin (compare **to Fig. 1**). These observations indicate a bimodal implication of actin polymerisation in N-cadherin-mediated adhesion that is necessary for the migration of cells displaying mild adhesion (Ncad cells), but prevents the migration of tightly adhering Ncad-FGFR cells, likely through the destabilisation of adhesions. Cell migration on N-cadherin thus requires an optimal adhesion that depends on the strength of the N-cadherin-F-actin mechanocoupling. To further support this hypothesis, we analysed the implication of myosin II, also contributing to the stabilisation of cadherin adhesion. Treatment with the myosin II inhibitor similarly blocked the migration of Ncad cells and stimulated the one of Ncad/FGFR cells (**Fig. S8**). Altogether, these data indicate that FGFR1 activity increases the coupling of N-cadherin complexes to the underlying cytoskeleton. The resulting strengthening of N-cadherin-mediated contacts contributes to the inhibition of cell migration on the N-cadherin substrate.

## Discussion

Although cadherin and FGFR dysfunctions are observed in cancer, their relation to cell migration and invasion remain unclear. N-cadherin facilitates either cell adhesion or cell migration, whereas FGFRs are either enhancers or repressors of cell migration. In light of reported cell type specific cadherin/tyrosine kinase growth factor receptor, E-cadherin and EGFR^44^ and VE-cadherin and VEGFR^6, 20^, functional interactions, a crosstalk between N-cadherin and FGFR has been proposed^4, 51, 64, 77^, although the mechanisms by which it may affect cell adhesion and migration remained unclear.

To mimic N-cadherin-dependent neural or cancer cells migration over neighbouring cells, we set up a model system of isolated N-cadherin or N-cadherin/FGFR1 expressing cells migrating on recombinant N-cadherin-coated lines. We describe here a complex interplay between N-cadherin engagement and FGFR1 activation positively regulating the strength of cell-cell adhesion and decreasing cell migration on N-cadherin, which occurs in the absence of added FGF. FGFR1 expression dramatically blocked N-cadherin-dependent single cell migration. This inhibition was associated to an increased cell spreading due to a strengthening of N-cadherin-mediated adhesion. FGFR1 led to the reinforcement of N-cadherin adhesion as demonstrated by the increased recruitment and stabilisation of junctional N-cadherin. We do not know whether FGFR1 regulates directly the “cis”- or “trans”-clustering of N-cadherin that may affect its stability at the plasma membrane^46^. However, this stabilisation was associated to an increased resistance of cell-cell contacts to calcium depletion, to an increase in the coupling of cadherin complexes to the actin treadmilling and to a rise in the mechanical strength of cell contacts. The rupture force of N-cadherin-mediated bead-cell contacts measured here was in the same range than those reported for doublets of N-cadherin expressing S180 cells (7.7 ± 1.4 nN)^9^. This rupture force was significantly increased by FGFR1 expression.

Altogether, our data strongly support the hypothesis that FGFR1 blocks cell migration on N-cadherin by strengthening N-cadherin adhesion. This behaviour is reminiscent of the reported biphasic relationship between cell migration of cells on fibronectin and the strength of integrin mediated cell-substratum adhesion^53, 55^. Cell migration is enhanced with increasing adhesion up to a threshold, above which further increases in adhesion acts to the detriment of migration. Accordingly, cells expressing only N-cadherin were poorly spread and supported cycles of both adhesion and deadhesion, allowing them to migrate and invert their polarity whereas cells expressing also FGFR1 remained tightly spread on the cadherin-coated substrate preventing migration. Pharmacological treatments altering actin polymerisation and actomyosin contraction, which destabilise cadherin adhesions^46^, blocked the migration of N-cadherin expressing cells, but stimulated the migration of cells expressing FGFR1. By analogy, the positive effect of FGFR signalling on the migration of neuronal cell growth cones^4, 77^ may be related to the intrinsic weak adhesion of neuronal cells.

The cellular responses reported in this study, including the regulation of N-cadherin stability at the plasma membrane and of the mechanocoupling between N-cadherin and actin require the receptor activation. Although we cannot exclude that FGFR1 could be activated by FGFs endogenously produced by HEK cells, these experiments have all been performed in the absence of exogenous FGF. Thus, we provide evidence that N-cadherin engagement by itself stimulates the activation of FGFR1, in agreement with previous observations made in neuronal cells^4, 72^ The presence of N-cadherin strongly decreases the mobility of junctional FGFR1 suggesting that the receptor was trapped in adhesion complexes. This process is N-cadherin specific, as junctional FGFR1 stabilisation was not observed with E-cadherin. We confirmed by co-immunoprecipitation, but also using purified proteins, that FGFR1 and N-cadherin interact through their extracellular domain. This may be essential for FGFR1 activation; decreasing the mobility of the receptor and/or increasing its local density at cellcell contacts may stimulate its dimerisation and cross phosphorylation.

The sustained activation of FGFR1 significantly increased N-cadherin levels at the plasma membrane. The level of expression of the p120 catenin has been reported to stabilise junctional cadherin by preventing their internalisation^12, 24, 50^. Accordingly, we found that a mutant of N-cadherin impaired for its binding to p120 was not stabilised at cell-cell junctions in FGFR1 expressing cells. Cells expressing this mutated cadherin together with FGFR1 spread poorly on N-cadherin and migrated at high speed. Thus, we propose that p120 is involved in the regulation of N-cadherin stabilisation at cell adhesion sites by FGFR1. It has been reported that the phosphorylation of this catenin may induce its dissociation from cadherin allowing endocytosis of the latter^24^. Thus, the negative effect of FGFR1 on p120 phosphorylation might be a relay to stabilise junctional N-cadherin. However, no changes were observed in the cellular levels of p120 tyrosine phosphorylation upon FGFR1 expression, in agreement with a previous study reporting that mutation of Y228 and other prominent Src-associated p120 phosphorylation sites did not noticeably reduce the ability of E-cadherin to assemble AJs^41^. Alternatively, p120-dependent endocytosis of N-cadherin upon FGFR expression may rely on phosphorylation on serine residues, which have been reported to regulate p120 functions^14, 33^. Alternatively, the increased stability of N-cadherin at cell-cell contact may depend by the reported regulatory action of p120 on Rho GTPases activity^1, 52^.

The kinase activity of the receptor was only partially involved in the inhibition of N-cadherin endocytosis. Moreover, the N-cadherin mutant impaired for p120 binding displayed a reduce association to FGFR1, indicating that the regulation of N-cadherin availability at the plasma membrane by FGFR1 involves additional pathways. Accordingly, we unravelled an involvement of Src in the cellular response to FGFR1 expression, which is unrelated to p120. FGFR1 increased the amount of activated Src associated to N-cadherin immunocomplexes and blocking pharmacologically Src activity reverted the effect of FGFR1 on cell spreading and migration. The inhibition of Src had a blocking effect on the stabilisation of junctional N-cadherin induced by FGFR1, indicating that Src is also involved in the mechanocoupling between N-cadherin complexes and actomyosin. Altogether, N-cadherin stability at the plasma membrane inversely correlates with the migratory properties of the cells on N-cadherin substrates. It is important here to recall that, in the case of the radial migration of cortical neurons *in vivo*, efficient migration on radial glia requires an active recycling of N-cadherin in neurons^17, 25, 29^. In this system, both the blockade of N-cadherin recycling and N-cadherin overexpression induced abnormal stabilisation of cell-cell contacts and impaired cell migration.

Taken together, these data reveal the existence of a pathway controlled by FGFR1 and N-cadherin and regulating N-cadherin-dependent cell-cell adhesion and cell migration. FGFR1 and N-cadherin are co-recruited and co-stabilised at the cell-cell adhesions. This leads to sustained activation of FGFR1, which in turn promotes N-cadherin accumulation at the plasma membrane, strengthens N-cadherin mediated cell-cell contacts and N-cadherin mechanocoupling to actin. Adhesion between migrating cells and N-cadherin-expressing cellular substrates is increased therefore decreasing cell migration. This mechanism could be used by cancer cells to engraft to the vessel wall or the host tissue. In less adherent cells, such as neurons or for cancer cells in other locations or considering different type of cancer cells, depending on the level of expression of N-cadherin and the dynamics of the actomyosin cytoskeleton, the same pathway may promote cell migration or cell anchoring.

## Materials and Methods

### Plasmid constructions

The construct encoding GFP-FGFR1 was constructed as described in supplementary methods using the mousse fgfr1-IIIc full sequence (gift from D. Ornitz, University of Washington).

### Cell culture and transfection

HEK 293 (HEK), C2C12, 1205Lu and U2OS cells were grown, transfected and selected, as described in supplementary methods.

### Drug treatments

The FGFR kinase activity inhibitor, PD173074 (Sigma, 20 nM final concentration), and the Src family proteins inhibitor, PP2 (Abcam, 100 nM final concentration), were added in the medium 30 minutes prior the beginning and maintain throughout the experiments. Hydroxy-dynasore (Sigma, 100 nM final concentration) was incubated for 1 hour.

### FGFR activation and Ca^2+^ switch assay

C2C12 cells cultures were starved in serum-free medium 24 hours and then treated for 5 minutes with 1ng/ml of FGF2^30^. Alternatively, starved cultures were first treated with 4 mM EGTA for 20 minutes and then washed and incubated in the presence of 5 mM of Ca^2+^.

### Ncad-Fc line guided cell migration

Guided cell migration on 10 μm wide N-cadherin coated lines was performed as described in supplemental methods, in the absence of exogenously added FGF.

### Fluorescence Recovery after Photobleaching

Dual wavelength fluorescence recoveries after photobleaching (FRAP) was performed at 37°C (see supplemental methods) and analysed as reported previously^63^.

### Magnetic tweezers

A homemade magnetic tweezer was the source of the magnetic field gradient used to pull Ncad-Fc coated paramagnetic microbeads attached to the cells (see supplemental methods). For the measurement of the rupture force of N-cadherin-mediated bead-cell contacts, Ncad or Ncad/FGFR cells seeded on 10 μm-width fibronectin coated lines were incubated with 4.5 μm magnetic Ncad-Fc coated beads for 30 minutes, then unbound beads were washed out. The magnetic microneedle was approached while cells and the moving tip were imaged in phase contrast (every 10 milliseconds during 2 minutes).

## Abbreviations

FGF: Fibroblast Growth Factor
FGFR: Fibroblast Growth Factor Receptor
FP: Fluorescent Protein
FN: Fibronectin
FRAP: Fluorescence Recovery After Photobleaching
GFP: Green Fluorescent Protein
Ncad: N-cadherin
PBS: Phosphate Buffer Saline
RTK: Receptor Tyrosine Kinase
SEM: Standard Error of the Mean

## Acknowledgement

This work was supported by grants from CNRS, ARC foundation (contract number: PJA 20151203185), Human Frontier Science Program (HFSP grant RPG0040/2012), European Research Council under the European Union’s Seventh Framework Program (FP7/2007-2013) / ERC grant agreements n° 617233 (BL), Agence Nationale de la Recherche (ANR 2010 Blan1515) and NUS-USPC exchange program. TN was supported by a HFSP grant RPG0040/2012, then by FRM (Fondation pour la Recherche Médicale) and Labex WhoAmI. We would like also to thank all present and past member of the Cell Adhesion & Mechanics lab at the Institute Jacques Monod for constant support and exchange. We thank Region Ile de France (E539) and Ligue contre le Cancer (R03/75-79) for the acquisition of the equipment. We thank C. Murade and M. Yao for held with magnetic tweezer experiment. C.M. was supported by Fondation pour la Recherche Médicale (FDT20150532600), L.D. by the Association pour la Recherche contre le Cancer (Fondation ARC, P2009 CDD POST-DOC) and D.G.F by North West Cancer and the Cancer and Polio Research Fund. We thank O.Thoumine and R. Horwitz for their kind gift of NcadAAA and FGFR1 encoding plasmids, respectively. We acknowledge the ImagoSeine core facility of the Institut Jacques Monod, member of IBiSA and France-BioImaging (ANR-10-INBS-04) infrastructures.

