## Supplementary Figures and Methods for "Enhanced cell-cell contact stability and decreased N-cadherin-mediated migration upon Fibroblast Growth Factor Receptor-N-cadherin cross-talk"

### Figure S1

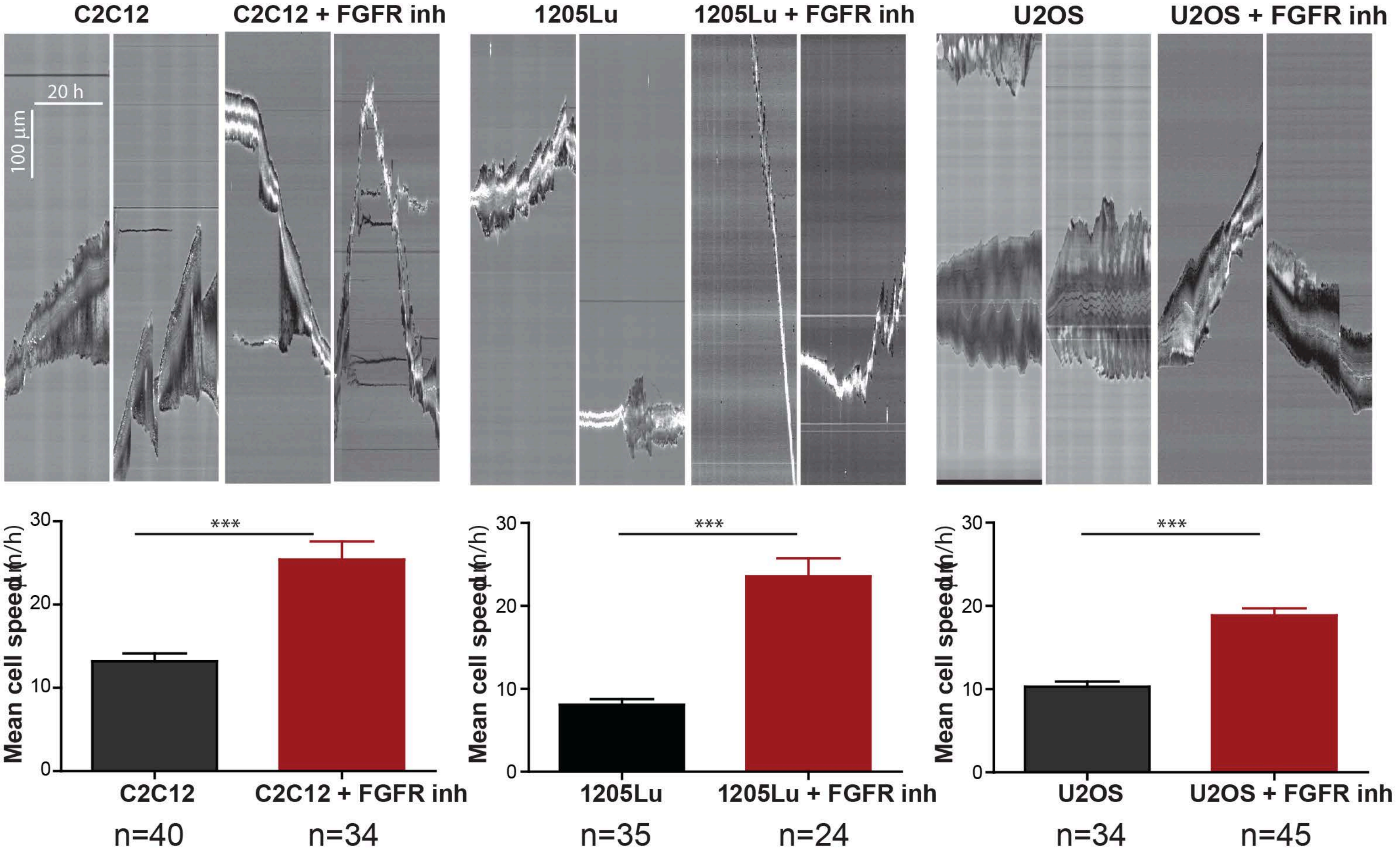

C2C12
C2C12 + FGFR inh
1205Lu
1205Lu + FGFR inh
U2OS
U2OS + FGFR inh
20 h
100 µm
Mean cell speed (µm/h)
30
20
10
0
***
C2C12
C2C12 + FGFR inh
1205Lu
1205Lu + FGFR inh
U2OS
U2OS + FGFR inh
n=40
n=34
n=35
n=24
n=34
n=45

### Figure S2

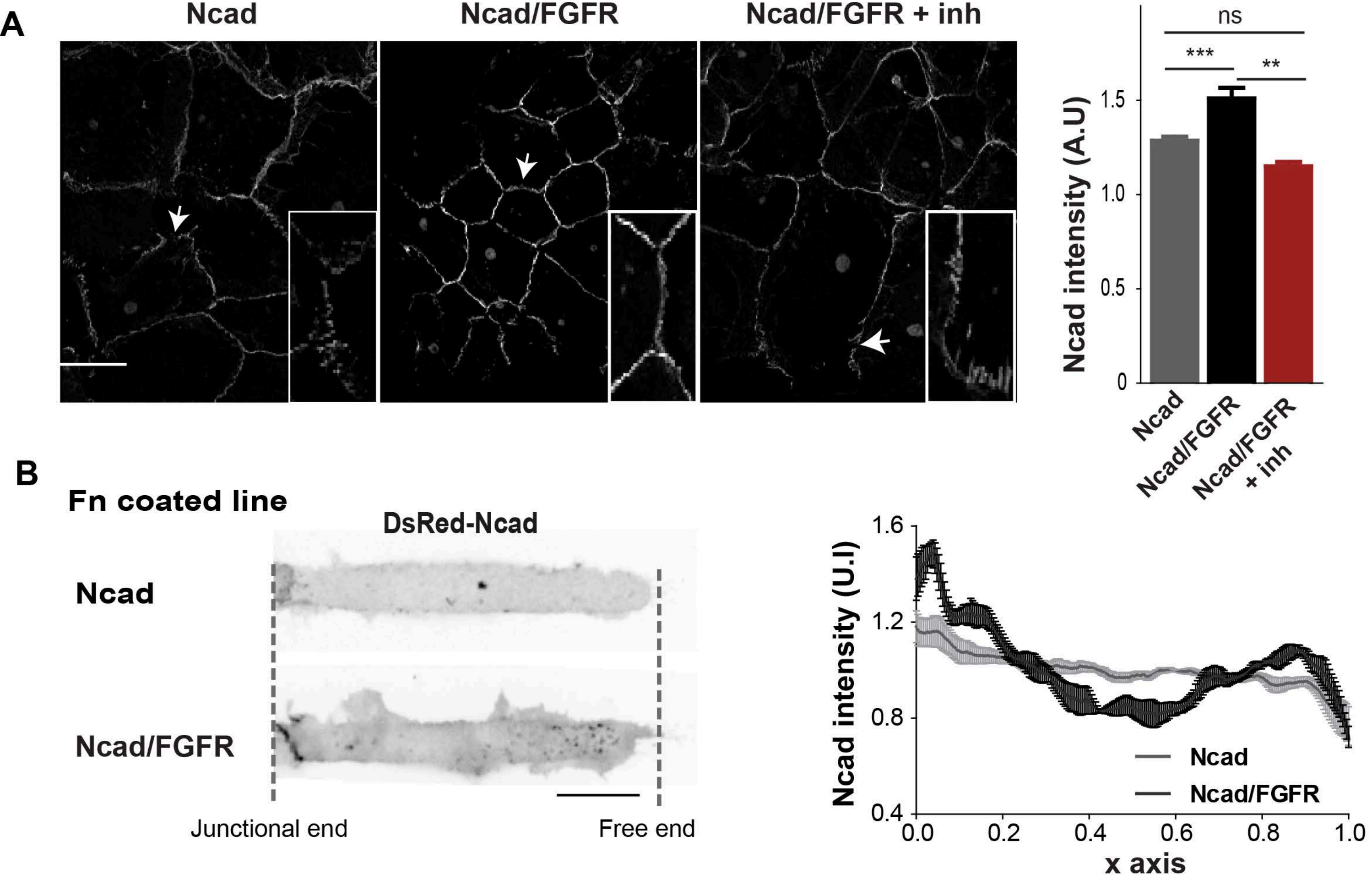
A
Ncad
Ncad/FGFR
Ncad/FGFR + inh
ns
***
**
Ncad intensity (A.U)
1.5
1.0
0.5
0
Ncad
Ncad/FGFR
Ncad/FGFR + inh
B
Fn coated line
DsRed-Ncad
Ncad
Ncad/FGFR
Junctional end
Free end
Ncad intensity (U.I)
1.6
1.2
0.8
0.4
0.0
0.2
0.4
0.6
0.8
1.0
x axis
Ncad
Ncad/FGFR

### Figure S3

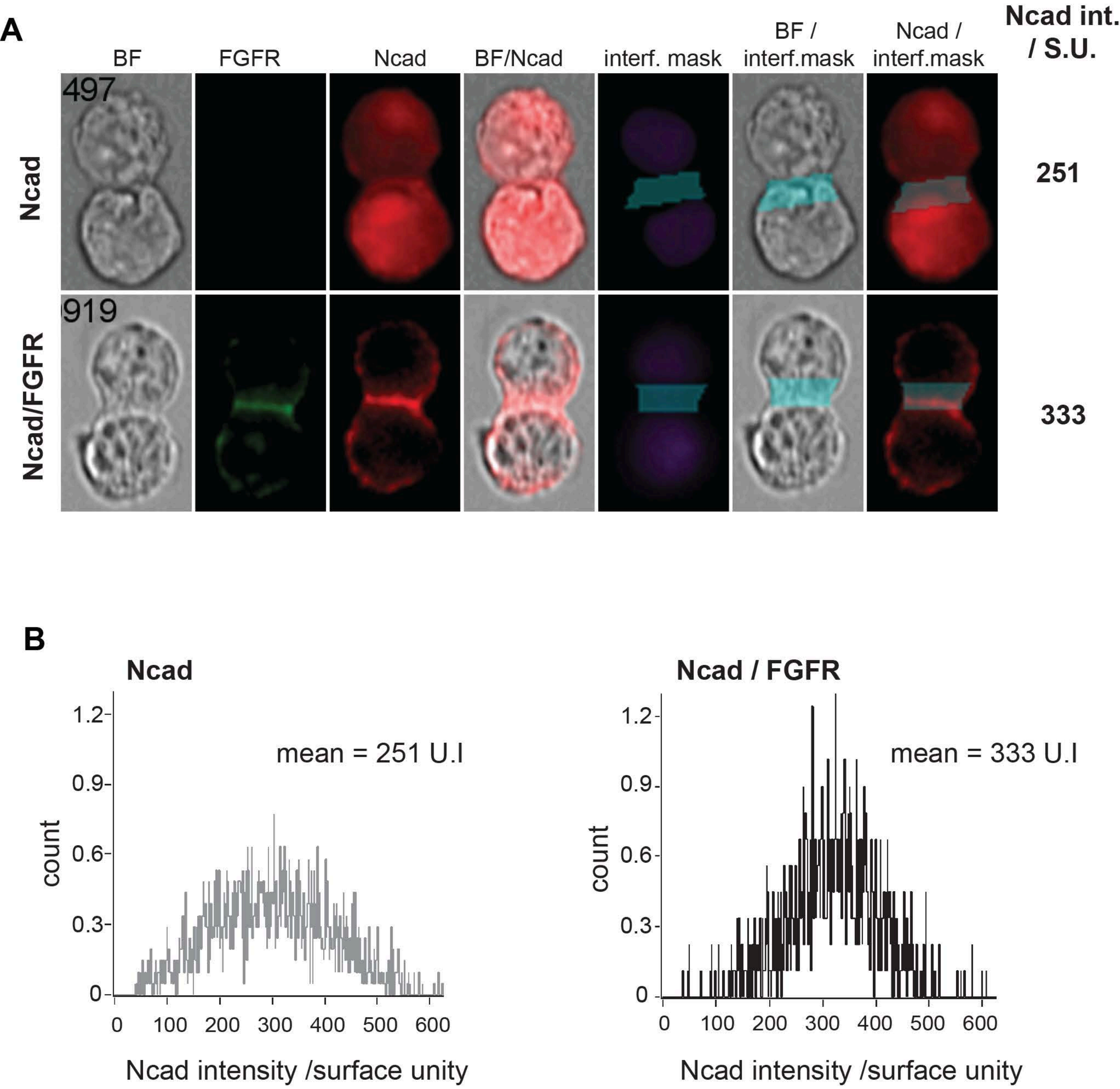
A
BF
FGFR
Ncad
BF/Ncad
interf. mask
BF / interf.mask
Ncad / interf.mask
Ncad int. / S.U.
Ncad
Ncad/FGFR
497
919
251
333
B
Ncad
Ncad / FGFR
mean = 251 U.I
mean = 333 U.I
count
Ncad intensity /surface unity
1.2
0.9
0.6
0.3
0
100
200
300
400
500
600

### Figure S4

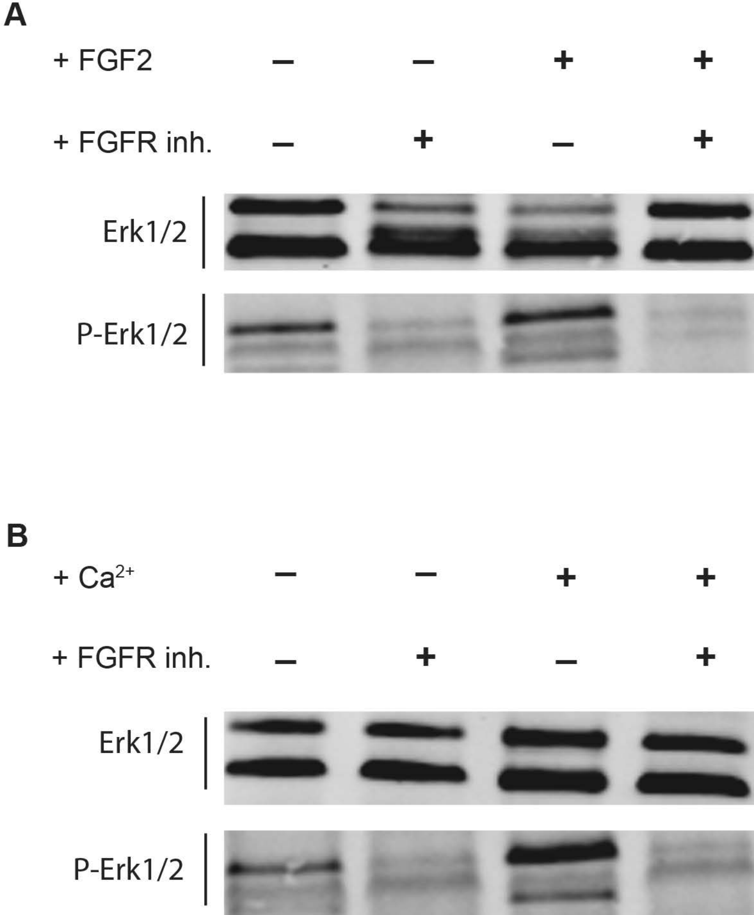

### Figure S5

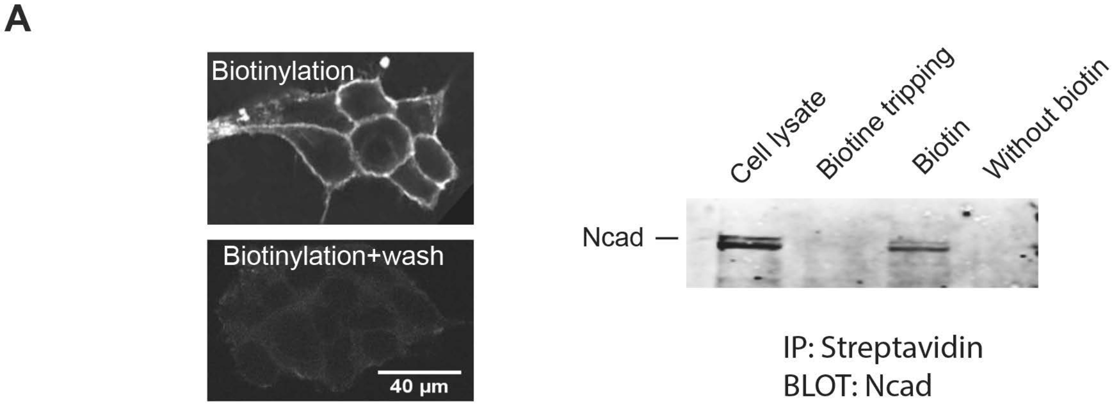
A
Biotinylation
Biotinylation+wash
40 µm
Cell lysate
Biotine tripping
Biotin
Without biotin
Ncad
IP: Streptavidin
BLOT: Ncad

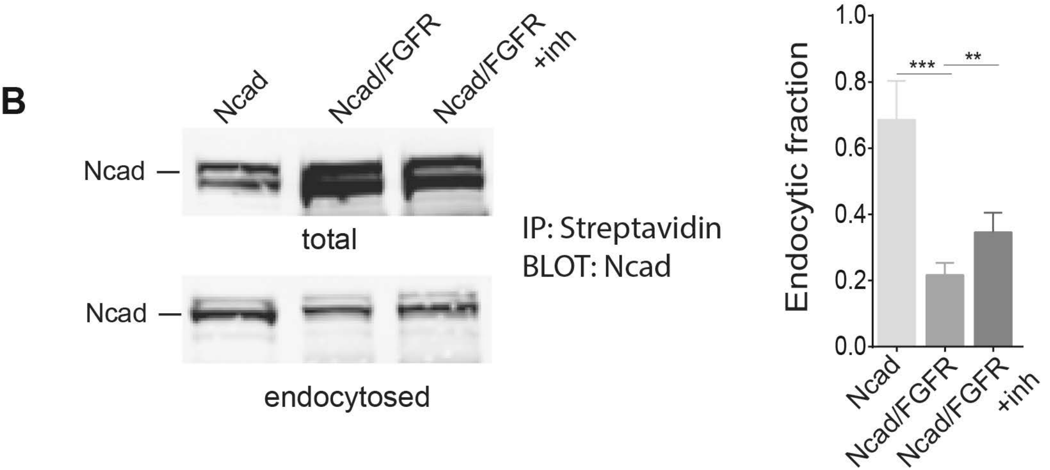
B
Ncad
Ncad/FGFR
Ncad/FGFR +inh
Ncad
total
Ncad
endocytosed
IP: Streptavidin
BLOT: Ncad
Endocytic fraction
1.0
0.8
0.6
0.4
0.2
0.0
***
**
Ncad
Ncad/FGFR
Ncad/FGFR +inh

### Figure S6

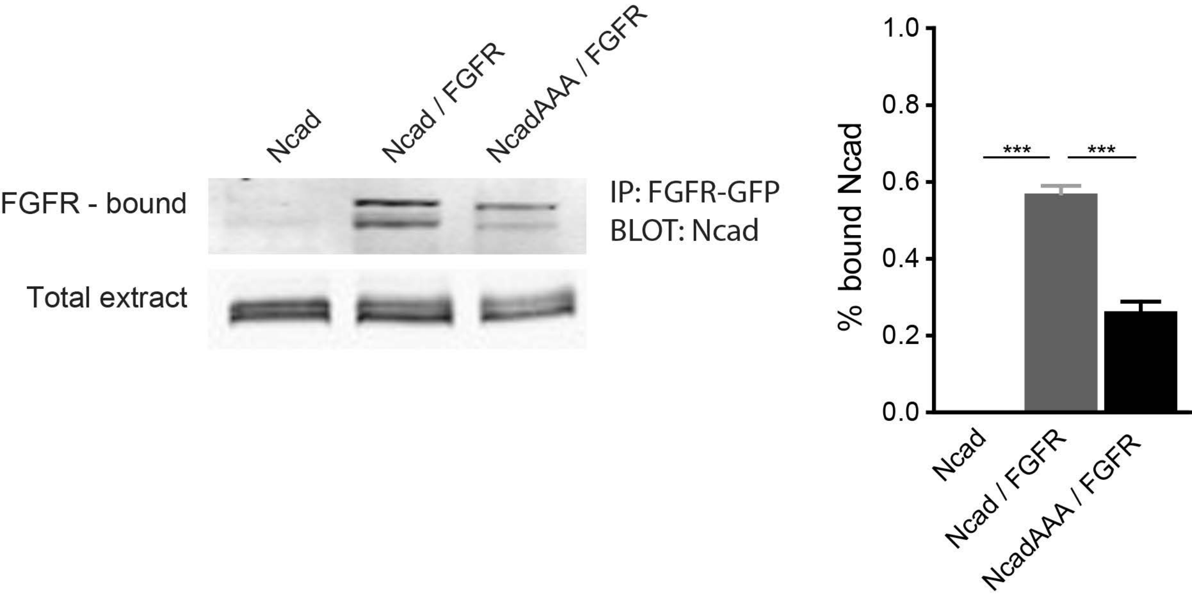

Ncad
Ncad / FGFR
NcadAAA / FGFR
FGFR - bound
Total extract
IP: FGFR-GFP
BLOT: Ncad
% bound Ncad
1.0
0.8
0.6
0.4
0.2
0.0
***
***
Ncad
Ncad / FGFR
NcadAAA / FGFR

### Figure S7

A

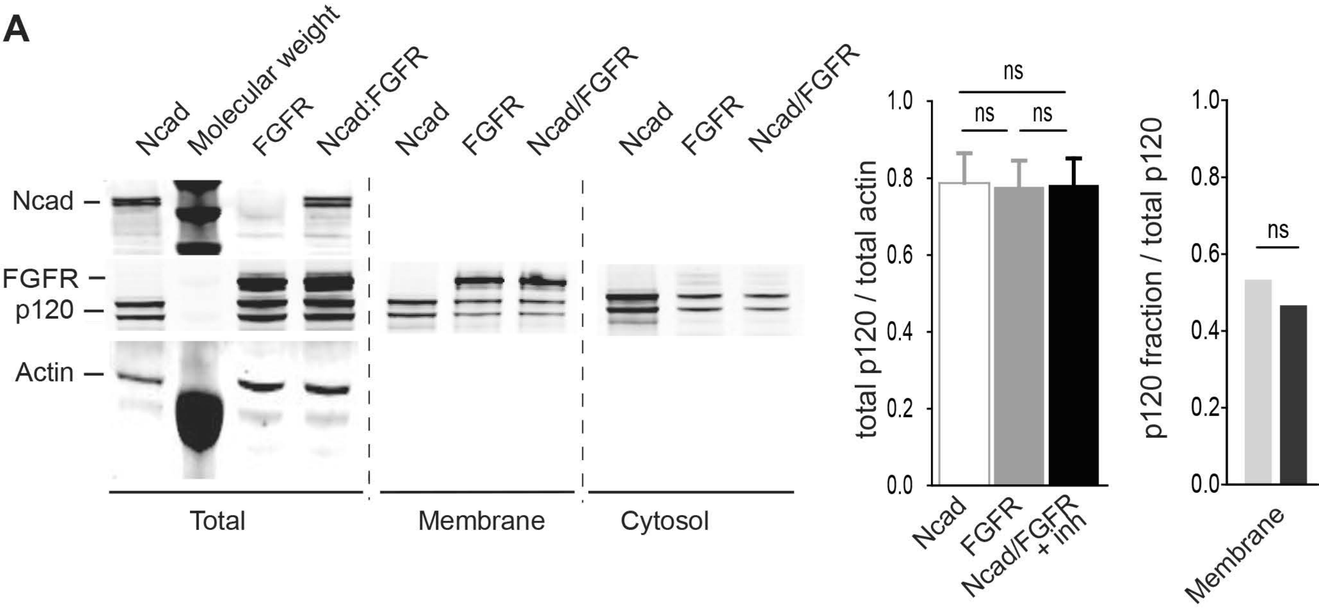
Ncad
Molecular weight
FGFR
Ncad:FGFR
Ncad
FGFR
Ncad/FGFR
Ncad
FGFR
Ncad/FGFR
Ncad
FGFR
p120
Actin
Total
Membrane
Cytosol
ns
ns
ns
total p120 / total actin
1.0
0.8
0.6
0.4
0.2
0.0
Ncad
FGFR
Ncad/FGFR + inh
p120 fraction / total p120
ns
Membrane

B

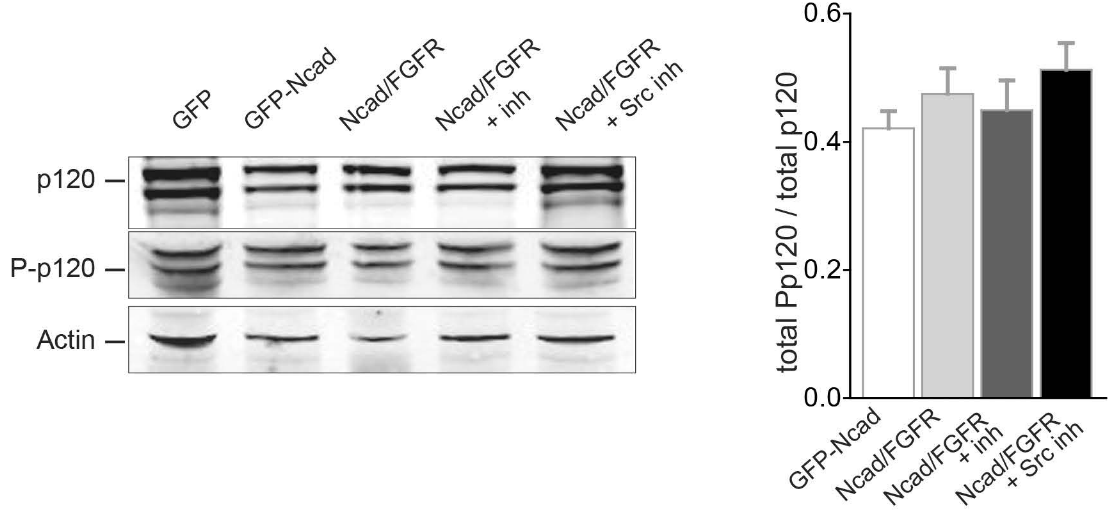
GFP
GFP-Ncad
Ncad/FGFR
Ncad/FGFR + inh
Ncad/FGFR + Src inh
p120
P-p120
Actin
total Pp120 / total p120
0.6
0.4
0.2
0.0
GFP-Ncad
Ncad/FGFR
Ncad/FGFR + inh
Ncad/FGFR + Src inh

### Figure S8

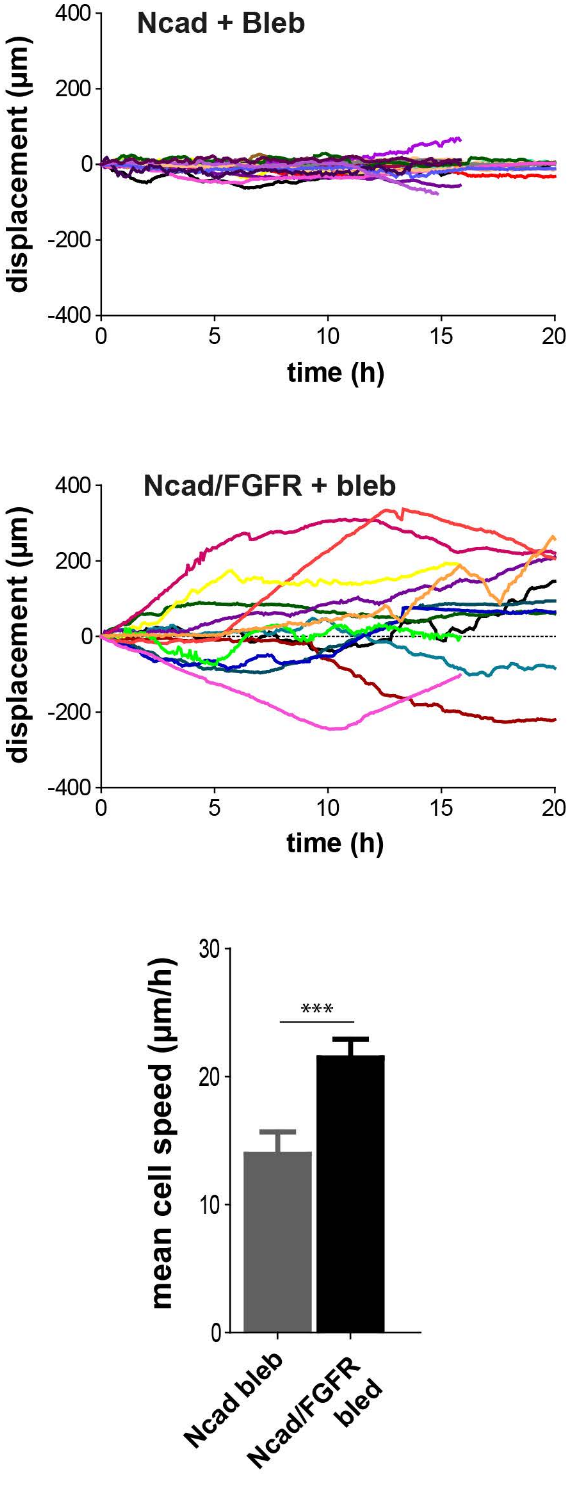
Ncad + Bleb
displacement (µm)
400
200
0
-200
-400
0
5
10
15
20
time (h)
Ncad/FGFR + bleb
displacement (µm)
400
200
0
-200
-400
0
5
10
15
20
time (h)
mean cell speed (µm/h)
30
20
10
0
***
Ncad bleb
Ncad/FGFR bled

#### Supplemental Figure Legends

**Figure S1: FGFR inhibitor treatment increases the migration of C2C12, 1205Lu and U2 OS cells on N-cadherin coated lines.** cells were seeded at low density on 10 µm-width Ncad-Fc coated lines in the absence or in the presence of FGFR inhibitor (+ FGFR inh) and imaged in phase contrast for 24 hours. Upper panels: Representative kymographs of the displacement of two cells for each condition. Lower panels: Histograms representing the mean cell speed C2C12, 1205Lu and U2OS cells, in the absence and in the presence of the inhibitor, respectively. *** $p \leq 0.0001$, ANOVA multi-comparison test, Newman-Keuls post-test).

**Figure S2: FGFR1 expression promotes N-cadherin recruitment and strengthens cell-cell contacts. (A)** DsRed-Ncad distribution in fixed monolayers of Ncad, Ncad/FGFR cells and Ncad/FGFR+inh cells grown overnight on glass coverslips in the presence of serum (fibronectin/vitronectin coating). Scale bar: 20 µm. Boxes show zoomed views of cell-cell contacts indicated by arrows. Cell-cell contacts in Ncad/FGFR cells appear straighter than those in Ncad cells and Ncad/FGFR+inh cells. Histograms on the right show DsRed-Ncad intensities measured at the cell-cell contacts in the three conditions thanks to Imaris. *** $p \leq 0.0001$; ** $p < 0.01$; ns: non-significant, ANOVA multi-comparison test, ($n \geq 50$). **(B)** Ncad and Ncad/FGFR cells were seeded on 10 µm width fibronectin-coated stripes and fixed after 2 hours. Left: cell doublets were imaged (scale bar: 20 µm). Right: the graphs show the mean distribution of DeRed-N-cad intensity along the cell width (z axis) normalised along the x axis of the cell with 0 value defined as the junctional edge of the cell and 1 value as the free edge, for Ncad and Ncad/FGFR cells (n = 30 and 27 doublets, respectively, SD = 0.177)).

**Figure S3: FGFR enhances the recruitment and the clustering of N-cadherin at cell-cell contacts. (A)** Analysis by flow cytometry imaging of Ncad recruitment at cell-cell interface in cell doublets. DsRed-Ncad recruitment at cell-cell contacts was quantified as the average normalised Ncad fluorescence intensity per surface unit in cell-cell areas (Ncad int./S.U.). FGFR expression significantly increased Ncad recruitment at cell-cell contacts. Data acquisition was performed for $1.5 \times 10^5$ cells for each condition and repeated 4 times. **(B)** Distribution plot of values obtained for the normalised Ncad fluorescence intensity per surface unit at cell-cell areas (Ncad int./S.U.) presented in Figure S2A.

**Figure S4: Enhanced N-cadherin engagement sustains activation of FGFR downstream pathway.** C2C12 cells were treated with FGF2 (5 nM, 15 minutes) or/and with FGFR inhibitor (10 nM, 1 hour) **(A);** or preincubated with EGTA (2 mM, 30 minutes), then switch back to medium containing 2 mM $Ca^{2+}$ for 10 minutes in the absence or in the presence of FGFR inhibitor **(B).** Cells were lysed and total extractions were subjected to electrophoresis and Western blotting using anti-Erk1/2 and anti-P-Erk1/2 antibodies.

**Figure S5: FGFR decreases N-cadherin internalisation. A.** Left: Fluorescent imaging of cell surface protein biotinylation; Cy5-conjugated streptavidin labelling of freshly biotinylated cells and of biotinylated cells following reducing wash. Surface biotinylated Ncad cells were lysed and incubated with streptavidin. Total lysate of biotinylated cells (Cell lysate), streptavidin precipitated lysates of biotinylated cells submitted immediately after to a reducing wash at 4°C (Biotin + stripping), of biotinylated cells (Biotin) and of non-biotinylated cells (Without biotin) were resolved by SDS-PAGE and immunoblotted using anti-Ncad antibodies. **B.** Analysis of N-cad endocytic fraction following cell surface biotinylation for Ncad, Ncad/FGFR and Ncad/FGFR + inh cells. Experiments were performed similarly than in Figure 6B except that the hydroxy-dynasore treatment was replaced by FGFR inhibitor treatment. (Left) Western blot detection of N-cad in anti-N-cadherin (total) and streptavidin pooled (endocytosed) proteins. (Right) Quantification of the N-cad endocytosed/total ratio obtained over 3 independent western blots. $^{**}p \leq 0.01$; $^{***} p \leq 0.001$; ns: non-significant, ANOVA multiple comparison test.

**Figure S6: p120 is involved in the binding of N-cadherin to FGFR.** Analysis of N-cad binding to FGFR for Ncad, Ncad/FGFR and NcadAAA/FGFR cells. Experiments were performed similarly than in Figure 4B. (Left) Western blot detection of N-cad using anti-N-cadherin antibodies in total cell lysate (total) and FGFR-GFP immunoprecipitated proteins (FGFR-bound). (Right) Quantification of the FGFR-bound/total ratio obtained over 3 independent western blots. $^{**}p \leq 0.01$; $^{***} p \leq 0.001$; ns: non-significant, ANOVA multiple comparison test.

**Figure S7: FGFR expression reduces the total amount and cytosolic fraction of N-cadherin and p120. (A)** FGFR, Ncad and Ncad/FGFR HEK cells were collected without detergent. Then total, cytosolic and membranous fractions were then separated as detailed in Material and Methods, the proteins extracted and immunoblotted for N-cad, FGFR and p120 for total extracts and FGFR and p120 only for the subcellular fractions. Histograms present the quantification of total level of p120 reported to actin, and membrane-associated p120 in membranous fractions versus total p120 cellular content, respectively. ns: non-significant, unpaired t test, n = 3 experiments. **(B)** Total protein extracts of GFP-HEK, Ncad, Ncad/FGFR and Ncad/FGFR cells treated with Src or FGFR inhibitor were separated and immunoblotted using anti-Pp120 and anti-p120 antibodies. Actin was used for protein loading control. The histogram shows the ratio of Pp120 over p120 in total extract, determined from the quantification of 3 independent immunoblots then converted to percentage.

**Figure S8: Effect of blebbistatin treatment on the migration of cells on N-cadherin lines.** Tracked displacements over 20 hours of Ncad (n = 12), Ncad/FGFR cells (n = 13). Histogram represents the mean cell speed.

**Video legends**

**Video 1** (1mn55s): Ncad, Ncad/FGFR, Ncad/FGFR + FGFR inh cells migration on Ncad coated lines.

**Video 2** (12s): Magnetic tweezer experiments on Ncad, Ncad/FGFR, Ncad/FGFR + FGFR inh cells.

**Video 3** (48s): Ncad trafficking at the leading edge of Ncad, Ncad/FGFR migrating cells.

**Video 4** (1mn55s): Ncad/FGFR and NcadAAA/FGFR cells migration on Ncad coated lines.

**Video 5** (1mn55s): Ncad/FGFR and Ncad/FGFR+ Src inhibitor migration on Ncad coated lines.

**Video 6** (43s): Measurement of Ncad/actin mechanocoupling in C2C12 cells.

**Video 7** (1mn55s): Ncad/FGFR and Ncad/FGFR+ CK666 migration cells on Ncad coated lines.

**Supplemental Material and Methods**

#### *Plasmid constructions*

The construct encoding GFP-FGFR1 (FGFR1 tagged with GFP at its carboxy-terminal extremity) was obtained using as a template pMIRB-FGFR1-Myc plasmid (gift from D. Ornitz, University of Washington), which encodes for the mousse fgfr1-IIIc full length sequence. By performing polymerase chain reactions (PCR) using sets of appropriate primers, (i) HindIII restriction site was introduced at 5' extremity (5'-GCGAAGCTTACCATGTGGGGCTGGAAGTGCC-3') while, (ii) stop codon was abolished, and AgeI restriction site was introduced at 3' extremity (5'-GCGACCGGTGGGCGCCGTTTGAGTCCACTGTT-3') of the FGFR1 encoding sequence. Resulting PCR product was subcloned into the pEGFP-N1 vector using the HindIII and AgeI restriction sites. The constructs encoding Flag-FGFR1 (FGFR1 tagged with a flag tag at its carboxy-terminal extremity) was obtained using also as a template pMIRB-FGFR1-Myc plasmid, by performing polymerase chain reactions (PCR). Using sets of appropriate primers, (i) NheI restriction site was introduced at 5' extremity (5'-GCGGCTAGCACCATGTGGGGCTGGAAGTGCC-3') while, (ii) a Flag tag encoding sequence and PmeI restriction site was introduced at 3' extremity (5'-CGCGTTTAAACTCACTTATCGTCGTCATCCTTGTAATCGGCGGCCCCGCGCCGTTT GAGTCCACTGTT-3') of the FGFR1 encoding sequence. Resulting PCR product was subcloned into the pcDNA3.1 hygro(-) vector using the NheI and PmeI restriction sites.

#### *Cell culture and transfection*

HEK 293 (Human Embryonic Kidney) and C2C12 mouse myoblastic, U2OS human osteosarcoma and 1205Lu human metastatic melanoma (gift of Lionel Larue, Inst. Curie, Paris) cells were grown in DMEM medium supplemented with 10% foetal bovine serum (FBS), 2 mM L-Glutamine, 100 IU of penicillin, 100 µg/mL streptomycin at 37°C in the presence of 5% $CO_2$. HEK cells were transiently electroporated with the plasmids encoding for dsRed-fused wild type N-cadherin (dsRed-Ncad), or N-cadherin 3A mutated in the p120 binding site (dsRed-NcadAAA)[1, 4] and/or with a plasmid coding for GFP- or Flag- FGFR1. Electroporation was performed with the Amaxa Cell Line Nucleofector (kit V, program X-032). To generate dsRed-Ncad, GFP-FGFR, dsRed-Ncad/GFP-FGFR dsR-Ncad/Flag-FGFR stable HEK cell lines, transfected cells were grown under a selection pressure of 200 µg/mL of Hygromycin B, 1 mg/mL of Geneticin or both. Drug resistant cells were then sorted out by

FACS (Influx 500 Cytopeia/BD-Biosciences) subcloned and further maintained with half of concentration of antibiotic pressure.

***Protein extraction and co-immunoprecipitation***

Proteins were extracted from 5-6 x $10^6$ cells. Cell cultures were rinsed in ice-cold PBS, detached with non-enzymatic detaching solution (Cell Dissociation Solution Non-enzymatic 1x, Sigma) and centrifuged at 200 rcf for 7 minutes. Cell pellets were suspended in lysis buffer (10 mM Tris HCl pH 7.5; 150 mM NaCl; 0.5 mM EDTA; 0.5% Triton) on ice. Cells were then passed slowly 10 times through a 26 gauge needle and left on ice for 1 hour with extensive pipetting every 10 minutes. Lysates were cleared by centrifugation at 20000 rcf for 10 min at 4°C. GFP-tagged proteins were then immunoprecipitated using magnetic GFP-Trap®-M beads accordingly to manufacturer instructions (Chromotek). Briefly, 25 µl of GFP-Trap®-M beads were washed 3 times with the wash buffer (10 mM Tris HCl pH 7.5, 150 mM NaCl, 0.5 mM EDTA). Fifteen µl of beads were added to 300 µl of protein extracts diluted in lysis buffer and tumbled end-over-end for 1 hour at 4°C. Beads were then magnetically separated, washed 3 times with the wash buffer, suspended in 100 µl 2x sample buffer plus reducing agent (NuPAGE, Invitrogen) and boiled at 95°C for 10 minutes to recover bound proteins. Proteins from the input and bound proteins were subjected to SDS-PAGE electrophoresis using Bis-Trisacrylamide 4%-12% NuPAGE gels (Invitrogen) then transferred on nitrocellulose membranes (0.45 µm, GE Healthcare). Membranes were blocked with 5% nonfat milk and incubated with the adequate primary antibodies and then with IRDye-coupled secondary antibodies (Rockland). The membranes were scanned using Odyssey Imaging System (LY-COR Biosciences).

***Surface biotinylation and endocytosis***

Cell cultures were chilled down to 4°C by three washes with cold PBS/$Mg^{2+}$/$Ca^{2+}$ then labelled with 1 mg/ml of NHS-SS biotin (Pierce) in PBS/$Mg^{2+}$/$Ca^{2+}$ for 12 minutes at 4°C under gentle rocking. Biotinylation was stopped by adding PBS/$Mg^{2+}$/$Ca^{2+}$, 50 mM glycine, 0.5% BSA at 4°C. Cells were washed twice in cold PBS/$Mg^{2+}$/$Ca^{2+}$ and lysed. Biotinylated plasma membrane proteins were then separated by precipitation with streptavidin-coated magnetic beads (Pierce).

For endocytosis analysis, biotinylated cells were returned at 37°C for 40 minutes. Cells were then chilled down with cold PBS/$Mg^{2+}$/$Ca^{2+}$ and bound biotin remaining at the cell surface was cleaved by incubating with 50 mM Glutathione in 75 mM NaCl, 10 mM EDTA, pH 7.5 for 15 minutes under gentle agitation. Cells were lysed and protein extracts were subjected to precipitation with streptavidin-coated beads or with GFP-Trap®-M beads (Chromotek).

***Cell Fractionation***

Cells cultures were washed with ice-cold PBS then scraped in 500 µl of detergent-free buffer **(**20 mM Tris-Cl pH 7.4, 100 mM NaCl, 2 mM MgOAc, 5 mM KCl, 10 mM GTP + protease inhibitor cocktail- both added fresh). Cells were dounced 35 times with a 26-gauge needle on ice and centrifuged at 450 rcf, 10mn. Supernatants were collected and centrifuged at 20000 rcf for 30 minutes. The supernatants, (cytosolic fraction) were collected. The pellets (membrane fraction) were rinsed with 1ml of lysis buffer then spun at 20000 rcf for 30 minutes. The pellets were re-suspended in 180 µl of detergent-free buffer supplemented with 1% NP-40 and incubated on ice for 1 hour while mixing every 10 minutes. The fractions were resolved by SDS-PAGE electrophoresis and immunoblotting.

***Fixed cell imaging***

Cell cultures were fixed at room temperature in PBS 4 % formaldehyde for 15 minutes. Preparations were then mounted in Mowiol, 90% glycerol. Images were acquired with a Leica TCS SP5 inverted confocal microscope AOBS tandem, equipped with a 63x oil objective (N.A=1.4), controlled by LAS AF (Leica System).

***Cell-cell contact disruption assay***

Glass or plastic surfaces were microcontact printed with silicon stamps bearing 200 µm fibronectin-coated squares as described in [6]. Cells were seeded on the patterned surfaces in the presence of 10 µg/mL mitomycin. After 1 h, unattached cells and mitomycin were washed out and preparations were returned to the incubator overnight. Preparation were processed for live image directly after addition of 5 mM of EGTA solution or fixed after 15 minutes of EGTA treatment. Live images were acquired at 20 x objective, every 30 seconds for 30 minutes under a controlled environment (37°C, 5% $CO_2$, type Inverted Olympus IX81, camera CoolSnap $HQ^2$) using MetaMorph. Fixed samples were acquired with the same microscope and camera, using 20 x and 60 x objectives.

***Cell doublets on fibronectin-coated line assay***

Stable dsR-Ncad and dsR-Ncad/Flag-FGFR HEK cell lines were transiently transfected with GFP-p120 or lifeact-GFP and seeded on fibronectine-coated-10 µm width lines[6], and let adhere for 1-2 hours. Samples were gently washed and returned to the incubator overnight. cell doublets were chosen to image in red-phenol-free DMEM supplemented with 10% serum every minutes during 1 hour under a controlled temperature and $CO_2$ environment (37°C, 5% $CO_2$, 40x oil objectives (N.A = 1.4), Spinning disk CSU22). Localisations of fluorescent proteins relative to the junction end were analysed using ImageJ for mask creating and Matlab for intensity calculation. All the fluorescence images were background-subtracted before quantification. The cells shape were detected by segmenting the fluorescence intensity image using Otsu method and converted into binary mask images with values outside the cell set to zero. The cell lengths were normalised to unity in the strip direction (x direction). For each individual cell, the fluorescence intensities within the cell mask along the x direction were averaged in the y direction (perpendicular to the strip direction) and projected in the x direction. The average intensity curves were normalised by the whole cell intensity and plotted against the normalised distance to the junction end. The average intensities in the x direction from multiple cells with the same experimental condition were calculated and an average curve was then created using Matlab function smooth by filtering with locally quadratic regression using a moving window of size 5. The overall behaviour of each group of multiple cells was then represented by one single curve.

***Ncad-Fc line guided cell migration***

Patterned silicon microcontact stamps bearing 10 µm width lines spaced of 70 µm were prepared by soft lithography according to a protocol derived from [6]. Patterned stamps were incubated with 1 µg/cm$^2$ anti-human IgG (Jackson Immunoresearch), pressed on non-culture treated petri dishes or on cleaned glass coverslip previously activated by deep UV (Jelight, 4 X 60W, 15 minutes). Microcontact printed surfaces were then passivated by incubation for 1 hour with 1% Pluronics F-127 (Sigma) diluted in distilled water, followed by 3 washes with PBS. Surfaces were incubated with 1µg/cm$^2$ hNcad-Fc (R&D) for 2 hours at room temperature then washed three times with PBS. Cells in culture were then dissociated on non-enzymatic detaching solution (Cell Dissociation Solution Non-enzymatic 1x, Sigma), seeded (10$^5$cells/200µl/cm$^2$) on these arrays of Ncad-Fc-coated lines and allowed to adhere for 1-2 hours in culture medium containing 1 µg/mL of mitomycin, before non-adhesive cells were gently washed off. Cells were imaged live or fixed 18 hours after seeding. For live imaging, Images were acquired with a 10 X objective, every 6 minutes during 24 hours under controlled environment (37° C, 5% $CO_2$, Biostation Nikon). Manual tracking of individual cells was performed with the MTrackJ plugin. Individual trajectories were positioned on an orthonormal axis with the coordinates of the cells at $t_0$ = (0:0). The displacements and mean cell speed were then extracted for each condition and plotted versus time and cell area, respectively.

***Fluorescence Recovery after Photobleaching***

Dual wavelength fluorescence recoveries after photobleaching (FRAP) was performed at 37°C on stably transfected Ncad, FGFR or Ncad/FGFR expressing cells, as well as on transiently transfected Ecad and Ecad/FGFR expressing cells. FRAP was performed using a Leica TCS SP5 confocal microscope equipped with a 40 X immersion objective (N.A=1.4) and carried out by setting the double scanning mode at 560 nm for dsRed and 480 nm for GFP and the image format to 256 x 256 pixels. After 3 prebleach scans (0.347 sec), a 20 x 40 µm ROI over the cell-cell contact was bleached with laser at full power by performing repeated scans. Recovery was recorded by imaging with low laser power every 0.347 sec (20 scans) then every 2 sec (20 scans) and finally every 10 sec (20 scans). The normalised recovery of fluorescence was expressed as a ratio of prebleach fluorescence after correction for photobleaching as reported previously[3]. Normalised fluorescence recovery in function of time curves were fitted with a one-term exponential equation using GraphPad Prism 5.01 software (one-phase decay non-linear regression functions5), allowing to extract a plateau value representing the fraction of diffusion-limited molecules (mobile fraction) and a recovery half-time ($t_{1/2}$) as a proxy of the apparent diffusion coefficient of diffusion-limited molecules[5].

***Flow Cytometry***

Cells were detached using non-enzymatic detaching solution (Cell Dissociation Solution Non-enzymatic 1x, Sigma), centrifuged at 200 rcf during 4 minutes, resuspended in culture medium and returned to the incubator for 10 minutes favouring moderate cell-cell adhesion in suspension. Cells were centrifuged again at 200 rcf for 4 minutes, fixed in 4% formaldehyde for 15 minutes, washed 3 times in PBS, incubated in the presence of 0.1% Dapi in PBS-BSA 0.1 % for 5 minutes, washed again then imaged under flow using ImageStream X (Amnis, Proteigene) set with the 405, 488, 560-nm laser and 480-560 filter. Data were analysed using the IDEAS software (Amnis, Proteigene) focusing on singlets for the quantifications of internal pool and on doublets for cell-cell accumulation of DsRed-Ncad fluorescence. For internal fluorescence quantification, regions corresponding to the total cell surface, the internal cell area and the cell membrane area were extracted from bright field images of singlets. Briefly, the morphology mask was applied to bright field images. Then, 4 pixels were evenly eroded from the border of the mask in order to exclude the cell membrane from the mask. The resulting mask was applied to the fluorescence channel. The internalisation feature was then applied to the final mask in order to calculate the internalisation score. Internalisation score per surface unit was defined as the ratio of internal fluorescence intensity per surface unit over the overall intensity per surface unit in the whole cell expressed on a logarithmic range.

The N-cadherin recruitment at the cell-cell interface was determined on doublet populations. Regions corresponding to the total cell surface and the cell-cell interface were extracted from bright field images and Dapi staining, respectively. The 4 pixels interface mask was determined as a region centred at the dimmest pixel between the 2 nuclei (Dapi). The interface mask was applied to the bright field channel to determine the surface area of the cell-cell contacts in the doublets, then to the fluorescence channel to count the intensity of Ncad staining at the cell-cell contact. Results were expressed as fluorescence intensity per surface unit.

***Production and biotinylation of soluble FGFR1 extracellular domain***

The soluble FGFR1 extracellular domain (FGFR1-ST) used here corresponds to amino acids 120 to 368 of the FGFR1IIIc comprising the acid box and immunoglobulin loops D2 and D3 with a poly-histidine-tail sequence followed by a thrombin cleavage site at the N-terminus and a Factor Xa cleavage site followed by a Strep-Tag II sequence at the C-terminus and is termed FGFR1-ST. This protein was produced in CHO cells and is heavily N-glycosylated [2]. For biotinylation, 2 µL of sodium periodate at 0.5 M freshly resuspended in water was added to 28 µL of purified FGFR1-ST at 16 µM in PBS supplemented by 0.01% of tween (PBS-T). The mixture was left incubated 1 hour in the dark. Excess sodium periodate was then removed using nanosep 30 kDa centrifugal device following the manufacturer recommendation and using PBS-T as exchange buffer. Purified periodated FGFR1-ST was recovered in 50 µL of PBS-T and 1.5 µL of biotin-LC-hydrazide at 135 mM in DMSO was then added. The reaction was left incubated overnight at RT and then kept 6 hours at 4°C. Excess biotin-Lc-Hydrazide was removed using nanosep 30 kDa centrifugal device following the manufacturer recommendation and PBS-T as exchange buffer. Biotinylated FGFR1-ST was recovered in 200 µL of PBS-T and kept at -20° c. Its concentration was estimated at 1.14 µM by measurement of the absorbance at 280 nm ($\varepsilon_{280\ nm}$ = 47424 mole$^{-1}$.L.cm$^{-1}$).

***Optical biosensor experiments***

Streptavidin (Sigma, 50 µl at 2.5 mg/mL) was immobilised on aminosilane surfaces using bissulfosuccinimidyl suberate ($BS^3$, Perbio, 1 mM) as the cross linker following the manufacturer's recommendations (NeoSensors, Sedgefield, UK). Surface was washed 5 times with 80 µL of Pi buffer (10 mM sodium phosphate buffer pH 7.2) and then incubated 3 min in 2M Tris-HCl, pH 8 to stop the cross-linking reaction. Three washes with 80 µL of Pi buffer and then of PBST were then performed before the addition of 25 µL of biotinylated FGFR1-ST at 1.14 µM. The reaction was left incubated for 4 to 6 hrs at RT and then overnight at 4°C. Cuvette was then cleaned by washing 3 times with 50 µL of PBST, NaCl 2 M in Pi buffer, PBST, HCl 20 mM and then PBST again.

Binding assays were carried out in TNC buffer (Tris-HCl20 mM pH 7.5, NaCl 150 mM, $CaCl_2$ 2 mM and Tween-20 0.02%). A single binding assay consisted of adding 0.5 to 6 µL of purified Ncad-Fc to a cuvette containing 19 to 24.5 µL or 24 to 29.5 µL of TNC. The association reaction was followed until binding was at least 90% of the calculated equilibrium value, usually between 150 s and 230 s. The cuvette was then washed three times with 50 µL TNC to initiate the dissociation of bound Ncad-Fc. Regeneration of the surface between each binding assay was performed by washing 3 times with 50 µL 2 M NaCl in Pi buffer, TNC, 20 mM HCl, and then TNC, which removed 98% to 100% of bound Ncad-Fc. Binding parameters were calculated using the non-linear curve fitting program FASTFit (NeoSensors). Each binding assay yielded four binding parameters, which are the slope of initial rate of association, the on-rate constant ($k_{on}$) and the extent of binding, all calculated from the association phase, and the off-rate constant ($k_{off}$, equivalent to the dissociation rate constant, $k_{diss}$), calculated from the dissociation phase. Biosensor experiments were carried out 4 times on 3 different FGFR1-ST-derivatised surfaces. The determination of binding kinetics in optical biosensors was prone to second phase binding sites at high concentration of Ncad-Fc. Thus, limiting amounts of ligand were immobilised on the sensor surface, whereas the slope of initial rate, $k_{on}$ and the extent of binding were only determined at low concentrations of ligate (3 different experiments), and $k_{off}$ was measured at higher concentrations of ligate (2 different experiments), to avoid steric hindrance and rebinding artefacts. A single site model was used to calculate all binding parameters. The dissociation constant ($K_D$) was calculated both from the ratio of the $k_{diss}$ and $k_{ass}$ and from the extent of binding, to provide an estimate of the self-consistency of the results.

***Magnetic tweezers***

The magnetic microneedle device was made of a 5 cm long stainless steel sewing needle glued to the top of permanent neodymium iron boron (NeFeB) surface surrounded by an aluminium rod. The montage was assembled on a micromanipulator (MP-285, Sutter Instrument) at a 30°C vertical angle, and the tip initially aligned at 600 µm from the centre of the observation field. The whole device was mounted on an inverted microscope (Olympus IX 81) equipped with a 40 x phase contrast air objective and a CCD camera (CoolSnap $HQ^2$) operating in the burst mode (frequency of 15 frames/s for 2 minutes). The micromanipulator allowed translational movement across all three axes with nanometer precision to position the magnetic field in the vicinity of beads. The distance between the tip of needle and detached bead was measured with imageJ.

The force applied to the bead is decreasing exponentially with the distance to the magnetic rod. To calibrate the magnetic force with is a function of the distance between the needle tip and the bead, 4.5 µm beads ware placed in a 100% polyethylene glycol solution (Mn 700, Sigma) at various distances of the needle and the bead motion was tracked by video microscopy. The instantaneous horizontal bead velocity ($v$) was extracted using ImageJ tracking. The force applied on the bead ($F$) was calculated respecting Stokes equation: $v = F/6\pi\eta r$, where $\eta$ is the dynamic fluid viscosity (for PEG Mn700: $\eta = 25$ Pa.s at 25°C) and $r$ is the radius of the bead. The calibration was performed ten times and the forces versus distance data were regressed to an exponential equation.


1 Bard L, Boscher C, Lambert M, Mege RM, Choquet D, Thoumine O. A molecular clutch between the actin flow and N-cadherin adhesions drives growth cone migration. J Neurosci 2008; 28: 5879-5890.

2 Duchesne L, Tissot B, Rudd TR, Dell A, Fernig DG. N-glycosylation of fibroblast growth factor receptor 1 regulates ligand and heparan sulfate co-receptor binding. J Biol Chem 2006; 281: 27178-27189.

3 Strale PO, Duchesne L, Peyret G, Montel L, Nguyen T, Png E *et al*. The formation of ordered nanoclusters controls cadherin anchoring to actin and cell-cell contact fluidity. J Cell Biol 2015; 210: 333-346.

4 Thoreson MA, Anastasiadis PZ, Daniel JM, Ireton RC, Wheelock MJ, Johnson KR *et al*. Selective uncoupling of p120(ctn) from E-cadherin disrupts strong adhesion. J Cell Biol 2000; 148: 189-202.

5 Thoumine O, Lambert M, Mege RM, Choquet D. Regulation of N-cadherin dynamics at neuronal contacts by ligand binding and cytoskeletal coupling. Mol Biol Cell 2006; 17: 862-875.

6 Vedula SR, Ravasio A, Anon E, Chen T, Peyret G, Ashraf M *et al*. Microfabricated environments to study collective cell behaviors. Methods Cell Biol 2014; 120: 235-252.
